# Multiscale Light Field Microscopy Platform for Multi-purpose Dynamic Volumetric Bioimaging

**DOI:** 10.1101/2024.09.30.615205

**Authors:** Yangyang Bai, Matt Jones, Lauro Sebastian Ojeda, Janielle Cuala, Lynne Cherchia, Senta K. Georgia, Scott E. Fraser, Thai V. Truong

## Abstract

Light field microscopy (LFM) has emerged in recent years as a unique solution for fast, scan-free volumetric imaging of dynamic biological samples. This is achieved by using a microlens array in the detection path to record both the lateral and angular information of the light fields coming from the sample, capturing a 3-dimensional (3D) volume in a single 2-dimensional (2D) snapshot. In post-acquisition, the 3D sample volume is computationally reconstructed from the recorded 2D images, thus enabling unprecedented 3D capture speed, not limited by the typical constraint of physically scanning the focal plane over the sample volume. Up to date, most published LFM imaging setups have been specialized single-purpose platforms, optimized for a narrow performance window in field of view and resolution, thus hampering widespread adoption of LFM for biomedical research. Here, we present a versatile LFM platform for fast 3D imaging across multiple scales, enabling applications from cell to system-level biology on the same imaging setup. Our multiscale LFM is built as an add-on module to a conventional commercially available wide field microscope, and the various imaging applications, with different ranges of field of view and resolution, are achieved by simply switching between the standard microscope objectives available on the wide field microscope. Importantly, we provide an open-source end-to-end software package for calculating the system performance parameters, processing the experimentally measured point spread function, and light field 3D image reconstruction. We demonstrate the performance of our multiscale LFM platform through imaging the whole-brain activity map of seizures in larval zebrafish, calcium dynamics in ex vivo mouse pancreatic islets, and subcellular protein dynamics in cultured cells.

## I. INTRODUCTION

Biological processes inherently occur in three dimensions (3D), ranging from intra-cellular trafficking to whole brain neural network activity^1–3^. Observing these processes in their native 3D context is essential for gaining a comprehensive understanding of how living systems develop, and how they function in health and malfunction in disease^4–6^. Volumetric microscopy techniques have greatly advanced our understanding of dynamic biological systems in 3D, yet challenges remain in capturing fast changing biological events at the relevant spatial-temporal resolution and coverage. Conventional volumetric imaging systems, such as confocal microscopy or Light Sheet Microscopy (LSM), acquire 3D volumetric images by scanning, either point by point or plane by plane across the entire volume^7,8^. While this strategy renders high resolution images, it is inherently slow due to the sequential scanning nature. Light Field Microscopy (LFM), on the other hand, obtains 3D volume in a single 2D snapshot without scanning^9–12^. This is achieved by inserting a micro-lens array (MLA) in the detection light path, recording synchronously both the spatial and angular information of the 3D sample’s light field onto the 2D camera. The 3D information is then reconstructed computationally in post-processing by solving the inverse problem with knowledge of the microscope optical train^13–15^. Because of the non-scanning nature, LFM has been demonstrated to reach a high volumetric imaging speed, reaching up to 100 Hz^16–18^, and opening up dynamic 3D imaging applications that are otherwise beyond the reach of scanning imaging modalities.

Currently, published LFM platforms have mainly been designed to work well for only single imaging scenarios^16–22^, where the focus is to maximize the imaging performance at a particular set of volume coverage and resolution. For example, particle tracking in cell biology was achieved in a setup with high imaging resolution (∼ 0.5 μm) and small depth of view (DOV) (∼ 6 μm)^19^, whereas whole-brain neural imaging in zebrafish larvae was implemented in platforms with larger DOV (∼ 250 μm) but lower resolution (≥ 2 μm)^16–18,21,22^. (Here, the DOV refers to the axial depth extent of the imaged volume where useful resolvable information can be collected and reconstructed.) These aforementioned examples were implemented with a specific set of hardware, lacking the flexibility to accommodate multiple imaging scenarios with diverse imaging performance requirements. One notable exception was the so-called “light field microscope eyepiece” which replaced one of the eyepieces of standard microscopes with a LFM module^23^. This approach enables convenient application of LFM to various imaging scenarios, though the eyepiece-mounted LFM module presents optomechanical constraints that could limit the overall performance of the system (see Discussion). and can be implemented as an add-on to a wide-field microscope using simple, off-the-shelf components. Another work described a scanning light field microscope (sLFM), which improved imaging resolution by scanning multiple axial positions over the volume and was implemented as an add-on to a wide field microscope using simple off-the-shelf optical components^24^. A detailed protocol for system design as well as a table listing different performance using different objectives and MLAs was provided. However, since the hardware system of the sLFM followed the design of the conventional LFM^9,13^, the numerical aperture (NA) of the MLA should match the NA of the primary objective, which means that each objective requires a matching MLA to achieve an ideal imaging performance (see Discussion). This significantly limits the versatility of the LFM for bio-imaging across different scales. We thus seek to expand further the effort to develop flexible LFM implementation both in hardware design and post-processing software pipeline to enable the widespread adoption of LFM by researchers who do not have expertise in optics, enlarging the impact that LFM would have on both basic and applied biomedical research.

In this work, we developed a versatile multiscale LFM platform that enables 3D bioimaging across a diverse set of imaging scenarios, in a flexible and user-friendly manner. We were inspired by the commercial Wide Field Microscopy (WFM) design where imaging at different scales can be achieved by simply choosing different microscope objectives housed in a rotating turret. Thus, we constructed our LFM based on a commercial WFM body, with an add-on LFM detection module connected to the standard camera port, and without any modification to the main microscope body (Fig. 1a 1, Supplementary Fig. 1). By taking advantage of the rotating turret, we reasoned that a wide range of bioimaging scenarios could be met by simply switching to a different primary objective without changing other hardware components in the LFM module (Table 1). This strategy marries the versatility and ease-of-use of the inverted microscope, a workhorse in many biomedical research labs, to the unique 3D imaging capability of LFM. We selected an off-the-shelf commercially-available MLA to achieve near optimal performance for all the objectives of interest (Fig. 1b). From the user’s perspective, our platform offers user-friendliness in several aspects. The sample is mounted in the same way as in an inverted WFM, and thus existing experimental workflows can be readily adopted for LFM imaging. For 3D reconstruction, we provide an open-sourced software package, along with a detailed, step-by-step procedure for acquiring and processing the experimentally measured point spread function (PSF) that is used in the reconstruction. To showcase the versatility of our LFM platform, we demonstrated its performance in three imaging scenarios that span a wide range of biologically relevant scales: characterizing whole-brain neuronal seizure in an intact zebrafish larva, recording calcium dynamic in an ex vivo mouse pancreatic islet, and observing subcellular protein dynamics in a cultured cell.

**Fig. 1.**
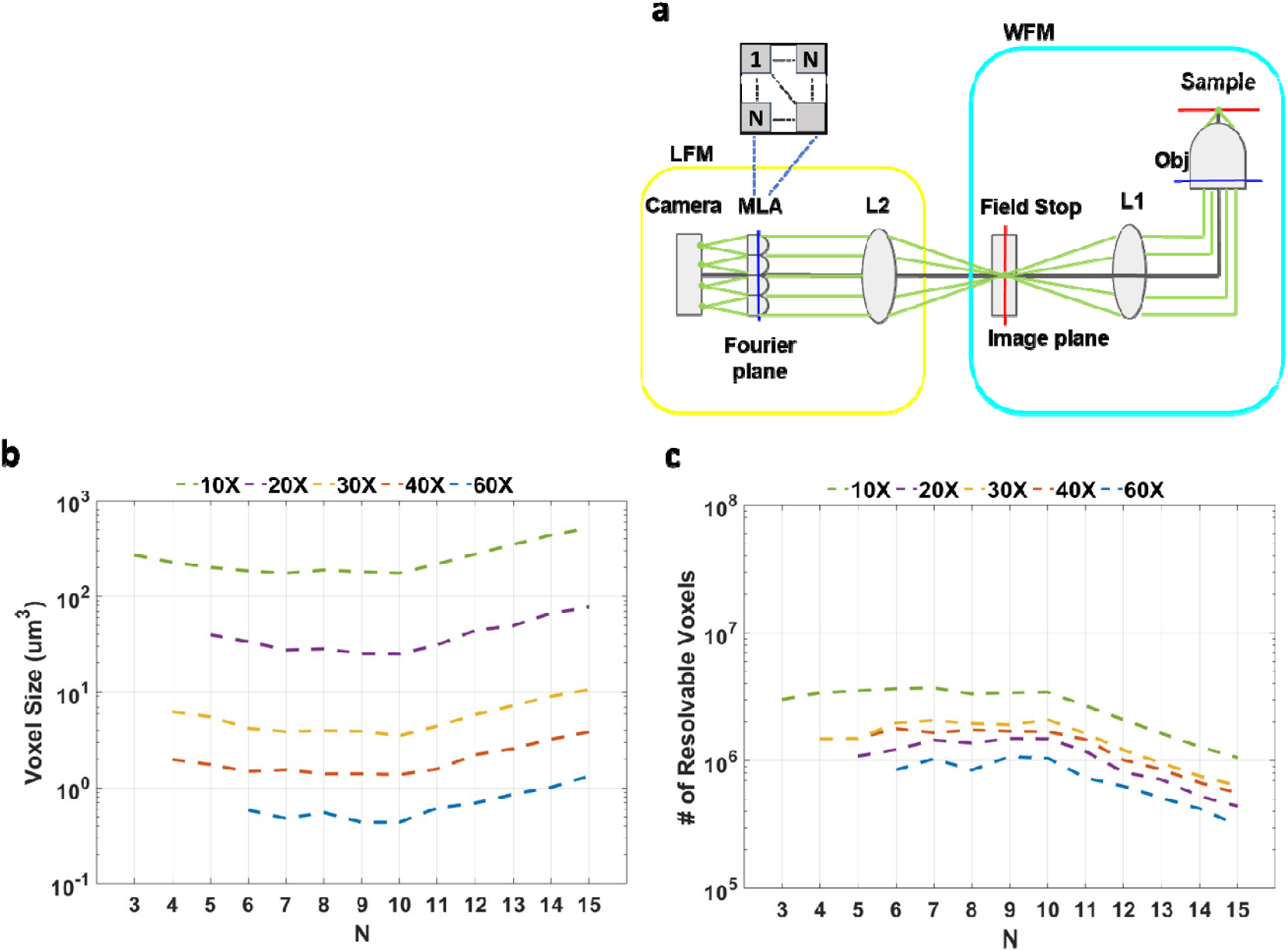
Design and performance of the multiscale LFM system. **a** Optical schematic of the system. Starting with the WFM body (Cyan), a field stop is placed at the standard image plane of the camera detection path. The LFM module (Yellow) consists of a relay lens L2, an MLA and a camera (Methods section). L2, in conjunction with L1 (the tube lens internal to the WFM), relays the back aperture (blue line) of the primary objective onto the MLA, which is placed at the back focal plane of L2. The camera is placed at the back focal plane of the lenslets of the MLA. Given a point source at the native object plane of WFM, the directional light ray bundle, depicted in Green, is resample at the MLA into multiple sub-aperture ray bundles and focused into multiple focal points behind each lenslet o to the camera, representing multiple angular views of the point source. Insert, schematic for an N × N square MLA. N is defined as the linear dimension of the square MLA. **b, c** Performance of all microscope objectives of interests, represented through, respectively, the single resolvable voxel size and number of resolvable voxels, as a function of N. The objectives vary over a wide range, from 10X 0.4 NA Air to 60X 1.35 NA Oil (Table 1).

**Table 1.**
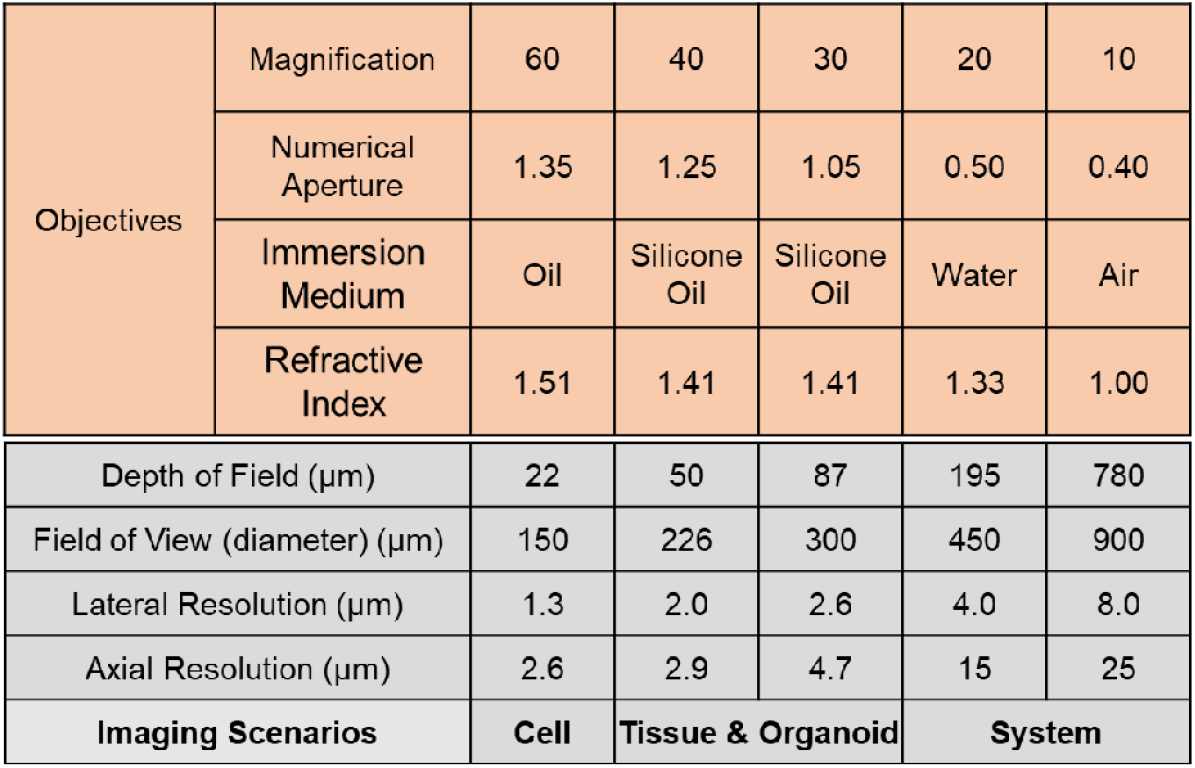
Estimated theoretical imaging performance of the multiscale LFM system. Top orange area lists all the objectives of interest for different bioimaging applications. The bottom gray area lists the corresponding imaging performance of the LFM module using each objective. The description of theoretical performance estimation was detailed in Supplementary Note 1. The corresponding wide field resolution is listed in Supplementary Table for all the objectives. As seen, with the range of standard microscope objectives depicted, the LFM module yields imaging performance that spans a wide range of spatial scales, enabling imaging across different imaging scenarios, from cellular to tissue to large-scale-system level.

## II RESULTS

### A. Overview of the multiscale LFM platform

Our multiscale LFM platform was constructed by adding a LFM module to the detection path of a commercial inverted WFM body (Methods, Fig. 1a, Supplementary Fig. 1). The rotating nosepiece (turret) integrated in the commercial microscope body holds multiple standard primary objectives that work directly with the LFM module (Supplementary Fig 1), allowing different sets of imaging performance to be obtained by simply switching between different primary objectives, without replacing hardware elements in the LFM arm. The LFM module is operated as an additional detection camera and is integrated into the existing image acquisition control of the WFM system.

In our platform, we implemented a recent version of LFM, termed Fourier Light Field Microscopy (FLFM)^10,11,14,15^, where an MLA was inserted at the Fourier plane of the detection optical light path (Fig. 1a). The MLA sub-divides the full aperture light field from the primary objective into multiple sub-aperture light fields. Each lenslet in the MLA focuses the corresponding sub-aperture light fields onto the image plane, generating multiple angular views of the 3D sample space behind each lenslet on the detection camera. This approach allows for recording, in the same 2D snapshot, both the lateral spatial information and the angular information of the light field. The original 3D sample space is recovered after image acquisition, using a deconvolution reconstruction algorithm and the experimentally measured point spread function (PSF) of the system^14,15^. Compared to the previous version of LFM where the MLA is placed at the image plane^9,13^, FLFM directly forms the angular views of a 3D sample behind each lenslet in the raw LFM image, allowing for immediate inspection and analysis on the raw LFM data without 3D reconstruction. Further, FLFM does not produce grid artifact at and near the native focal plane as observed in the previous LFM version, and has improved uniformity in lateral resolution across the DOV^11,14^. Importantly, the PSF of FLFM has lateral shift invariance, and thus the deconvolution operator in the 3D reconstruction can be performed in a parallelized fashion with GPU (Graphics Processing Unit)-accelerated Fast Fourier Transform (FFT), offering approximately 100 times faster reconstruction speed compared to the previous version of LFM^14^.

To design the hardware of the LFM module, we first matched the size of the MLA to the size of the camera chip so that all the angular views formed behind the MLA can be captured onto the camera, without needing an extra optical relay, and thus maintaining design simplicity. Since we used a common scientific Complementary Metal–Oxide– Semiconductor (sCMOS) camera from Hamamatsu (ORCA-Flash4.0) with chip size of 13 × 13 mm^2^, we chose a commercially-available square-shaped MLA with side length of 12 mm. The camera has the pixel size at 6.5 µm and pixel array dimension of 2048 × 2048.

We then considered two main design parameters: the focal length of lens L2 and the number of lenslets in the MLA (Fig. 1a). The focal length of L2, denoted here as f2, was chosen to be the same with the focal length of the tube lens in the WFM (L1). This formed a 1:1 relay ratio from the back aperture of the primary objective, i.e. the Fourier plane, to the back focal plane of L2 where the MLA was placed. With the simple 1:1 relay ration, the size of the optical aperture at the MLA is the same as the size of the objective back aperture. The chosen size of the MLA, 12 × 12 mm^2^, was found to be large enough to fully capture the back aperture sizes of the standard objectives used in our setup (Table 1, Supplementary Note 1). Next, we consider the number of lenslets in the MLA, which dictates the fundamental trade-off of the light field system: 3D imaging volume vs voxel resolution. For a square MLA, the number of lenslets can be represented by the linear dimension N defined as the number of lenslets on one side of the square MLA (Fig. 1a). From Supplementary Table 1 (and expanded in Supplementary Note 1), we observed that increasing N leads to a decreased Field of View (FOV) but increased DOV, as well as improved lateral and axial resolution. On the other hand, decreasing N results in an increased FOV but decreased DOV, and degraded lateral and axial resolution. These imaging performance parameters are changing in a competing fashion with respect to N. To optimize the trade-off, we evaluated how N affects two critical imaging performance metrics for 3D imaging: the size of the resolvable reconstructed single voxel, and the number of resolvable voxels within the captured volume. The equations to compute these two metrics are shown in Supplementary Note 1. The results are plotted in Figure 1b, c. At N ∼ 9, the resolvable voxel size reaches minimum, and the number of resolvable voxels reaches maximum, for all the standard primary microscope objectives. Considering off-the-shelf commercial availability, we chose an 8 × 8 square MLA for our system. The choice for the rest of the parameters in the LFM module is detailed in Supplementary Note 1. Altogether, our design choices, and using only commercially available components, results in a LFM imaging platform that could provide 3D imaging capability across a wide range of scales, from microns to sub-millimeters, covering cellular, tissue-level, as well as large-scale system-level imaging (Table 1).

The main limiting factor for resolution on our LFM system is the pixel size (6.5 µm), at which the full width at half maximum (FWHM) of the point spread function (PSF) is sampled at a rate below the Nyquist criterion. Reducing the pixel size increases the sampling rate, thereby pushing the resolution closer to the optical diffraction limit. For instance, with a camera pixel size of 2.2 µm, which was adopted in one previous work^10^, an ideal choice for N would be ∼7 (Supplementary Figure 5). The performances at pixel size 2.2 µm for three different N values are listed in Supplementary Table 6, 8, 10. Compared with performances obtained using our original 6.5 µm camera pixel size (Supplementary Table 5, 7, 9), the higher pixel sampling rate offered by smaller pixel size favors imaging applications requiring higher imaging resolution. However, smaller pixel sizes often come with trade-offs, such as increased cost, reduced signal-to-noise ratio (SNR), and lower frame rates. Cameras with larger pixels tend to be more cost-effective, offer better SNR, and support higher frame rates—features that are critical for capturing rapid dynamic processes at millisecond timescales. Given our emphasis on cost-efficiency and high-speed 3D imaging, we selected a camera with larger pixels to align with the practical goals of our light field imaging applications. Under either pixel size, N < 10 should be fine for a wide range of bioimaging applications, with better or worse performance for specific objectives. The final choice could be determined with a combination of imaging requirements, cost of manufacturing and commercial availability. Our contribution is focused on providing a metric and methodology with open-source design software to help people to calculate and decide for their own experiments.

The system performance was optimized with respect to N with fixed the focal length of MLA and relay lens (L_2_). The theoretical global optimization for FOV, DOV, R_xy_ and R_z_ with respect to N, focal length of MLA (f_ML_) and focal length of the relay lens (f_2_) is a complex non-linear multivariable optimization problem as shown in the equations in Supplementary Table 2. In this work, we fixed f_ML_ and f_2_ and mainly focused on optimizing N (or pitch of the lens-let) to make this problem more approachable for a broader audience who are not specialized in optical design. To further approach this problem, we use 30x 1.05NA Silicone objective as an example and plotted the system throughput (total number of resolvable voxels) and voxel resolution (Supplementary Figure 7) with respect to f_ML_ and f_2_ for each N to demonstrate the influence of these two parameters. In general, larger f_2_ and f_ML_ increases system throughput and degrade voxel resolution. An ideal choice for f_ML_ and f_2_ would be a balance between these two trade-offs. For our setup at N = 8 in Supplementary Figure 7e, the ideal f_2_ and f_ML_ for the number of Voxels and the voxel size are marked in red and cyan crosses respectively, which represents the greatest number of resolvable voxels (red cross) and the least single voxel size (cyan cross). Ideally, a good choice of f_2_ and f_ML_ would be in the middle area of the two crosses. Our choice of f_2_ and f_ML_ are marked in white cross. For the 30x objective, it is not an exact global optimal. However, it is important to note that our goal is not to achieve absolute optimal performance, which typically requires multiple customized optical components that could be difficult to obtain. Instead, we present a streamlined and effective design pipeline that can be readily implemented using commercially available components.

### B. 3D reconstruction pipeline with experimentally measured point spread function (PSF)

Richardson-Lucy (RL) deconvolution was used in our pipeline for LFM 3D reconstruction ^25,26^. The algorithm uses the PSF of the system and the raw LFM image to iteratively update voxel values in the 3D sample space. We obtained the PSF experimentally by imaging a sub-resolution-limit bead over the entire DOV. In contrast to a theoretical PSF, the experimental PSF captures the hardware imperfections deviating from the theoretical design, and thus enhances the accuracy of the 3D reconstruction^22^. For each axial plane, the PSF profile is shift invariant and the bead only needs to be imaged once at the center position^14^. An end-to-end software package written in MATLAB for LFM design, experimental PSF processing and LFM 3D reconstruction is provided openly on GitHub (Supplementary Note 2), along with video tutorials on YouTube. This feature is missing in the literature.

A unique feature of FLFM is that, in each angular view, the centroid of the PSF changes between different axial planes, encoding both lateral and axial information in one image. This is demonstrated in the Maximum-Intensity Projection (MIP) of the PSF with the axial position coded in different colors (Fig. 2a, b). In each angular view, different colors are distributed along a tilted line instead of overlapping over each other into a spot, as is observed for a conventional 2D imaging system, e.g. WFM. Due to the uniqueness of the FLFM PSF, none of the existing PSF extraction packages can be directly applied to the FLFM bead-imaging data. To extract the PSF from the raw bead image stack, we developed a PSF processing module that tracks the centroid of the PSF in each axial plane, ensuring an accurate and efficient cut-out of the PSF region (Supplementary Note 2). We package the PSF extraction module with the RL-deconvolution 3D reconstruction, together with additional image processing steps using Fiji ^27^, into an end-to-end open-source software pipeline, with a comprehensive user manual (Supplementary Software), to make our platform accessible to researchers without expert knowledge in LFM.

**Fig. 2.**
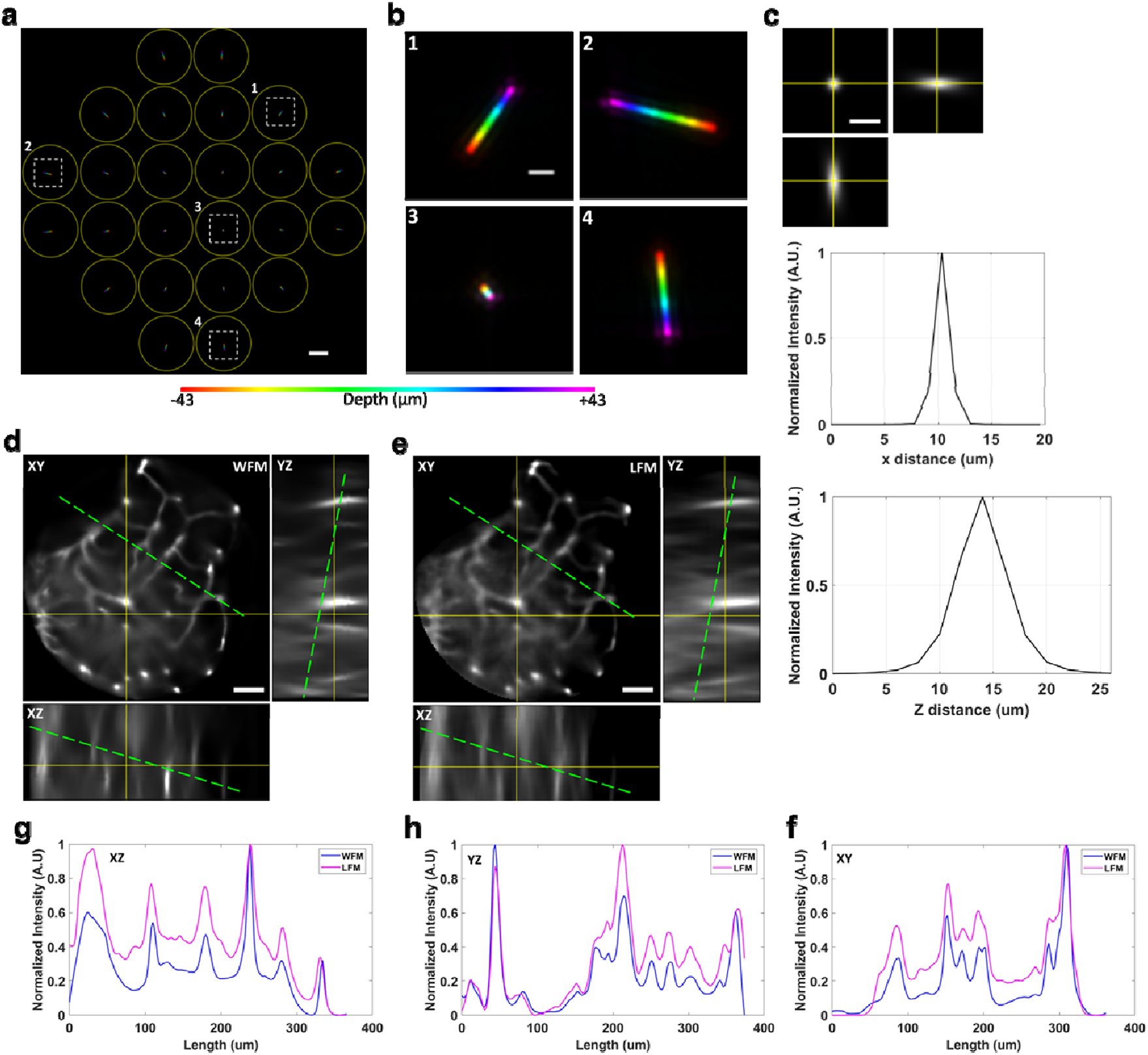
Resolution characterization and Ground Truth Imaging for the multiscale LFM. **a** Axial maximum-intensity projection of the PSF, taken with 30x 1.05 NA silicone oil immersion objective, where the axial z-position is color-coded over the DOV from -43 µm to 43 µm. The angular views behind each lenslet are circled out in yellow. Only the central views of the square MLA that are fully within the circular aperture of the objective are shown. **b** Zoomed-in views of dashed white boxes in **a.**Different angular views are seen as encoding z-depth (as colors) along lines oriented at different angles. **c** Lateral and axial image and line profiles of a reconstructed bead, recorded with the 30x 1.05 NA objective. See Supplementary Fig. 2 for the corresponding image and line profiles for the 60x 1.35NA and 10x 0.4NA objective. **d-e** Wide Field and Light Field imaging of Zebrafish brain vasculature. The zebrafish line used is kdrl:mCherry, labeling the cytoplasmic vasculature. The fish was imaged at 3 days post-fertilizatio (dpf), anesthetized with 0.016% tricaine and embedded in agarose for imaging. **d** Single plane Cross-sectional view of the 3D deconvolved WFM image stack. **e** Single plane Cross-sectional views of the 3D reconstructed FLFM image stack. **f** Line profiles of WFM and FLFM images along the green dotted line in the XY plane. **g**. Line profiles of WFM and FLFM images along the green dotted line in the XZ plane. **h**. Line profiles of WFM and FLFM images along the green dotted line in the YZ plane. A.U., Arbitrary Unit. Scale bars, (**a**) 100 µm, (**b**) 10 µm, (**c**) 5 µm. (**d,e**) 50 µm

The RL-deconvolution 3D reconstruction was implemented with GPU acceleration, carried out on a standard desktop workstation (Supplementary Note 2). Supplementary Table 3 listed the reconstruction times for all the datasets shown in sections D, E, F. Briefly, with our computing hardware, it takes ∼ 20 s to reconstruct a 2D (2048×2048) raw LFM image into a 3D volume. Supplementary Note 2 discusses practical ways to speed up the reconstruction process, such as down sampling the raw data (and sacrificing resolution), boosting the GPU processing power, or utilizing cloud computing services.

### C. Resolution characterization

We characterized the resolution of the LFM system by measuring the Full Width Half Maximum (FWHM) of the reconstructed single-bead images in both lateral and axial dimensions (Fig. 2, Supplementary Fig. 2). Since the PSF is acquired from imaging a sub resolution-limit single bead, each slice of the 3D PSF is equivalent to a LFM image of a single bead positioned at a different axial depth. Therefore, to reconstruct a single bead at different axial plane, we simply need to reconstruct several single slices from the PSF stack, below and above the native focal plane, using the full 3D PSF. We reconstructed five single-bead images, for each of the 10x, 20x, and 60x primary objectives. The lateral and axial FWHM values are found to be, respectively: 5.8 ± 0.1 µm, 31.0 ± 1.0 µm (10x 0.4NA); 1.8 ± 0.3 µm, 5.5 ± 0.2 µm (30x 1.05NA); 0.9 ± 0.04 µm, 3.2 ± 0.13 µm (60x 1.35NA) (Fig. 2c, Supplementary Fig. 2). For all three objectives, the FWHM measurements generally matched the theoretically-estimated lateral and axial resolution shown in Table 1. Notably, the lateral FWHM showed ∼ 30% improvement over the theoretically-estimated values (Fig. 2c). This is in fact not unexpected, as deconvolution has been shown to provide better lateral resolution performance than theoretical estimates based on wave-optics approximations^22^. Overall, the experimentally-measured imaging resolutions show that with the three objectives considered, the LFM provides the performance suitable for three different biological imaging scenarios: low-resolution large-volume system scale with the 10x 0.4NA, cellular-resolution medium-volume tissue scale with the 30x 1.05NA, and subcellular-resolution few-cells scale with the 60x 1.35NA. These imaging scenarios are demonstrated in the following sections.

### D. Imaging whole-brain seizures in zebrafish

We demonstrated the ability of our multiscale LFM in capturing fast biological system-level dynamics through imaging whole-brain seizures in larval zebrafish. Seizures are large-scale whole-brain neuronal population activity that unfolds within several seconds. Capturing the whole-brain seizure events in the zebrafish requires both fast imaging speed as well as a large imaging volume. Using the 10x objective (Table 1), we imaged the whole brain volume of ∼ 500 × 800 × 300 (XYZ) μm^3^ at 30 volumes/s (Supplementary Video 1), significantly faster than the state-of-the-art seizure imaging achieved by LSM operated at ∼ 5 volumes/s^28^. Even though seizure events in zebrafish generally last a few seconds to minutes ^29,30^, the high volumetric imaging speed allows for capturing of the fast dynamics within each event, especially during the rapid onset window. By analyzing the time series data from a cropped-out single lenslet image, which has data size at two orders of magnitude smaller than the full dataset, we could identify the existence and precise timing of the seizure event - this is an important capability to facilitate experimental trouble-shooting and efficient analysis of the large datasets. In this experiment, we reconstructed a 14-second movie for one representative seizure event (Fig. 3a, b). The averaged intensity over the reconstructed whole brain provides a coarse description of the seizure temporal profile (Fig. 3c). To highlight the propagation of brain activity across the brain during the seizure onset, we performed a temporal difference analysis on the 3D reconstructed movie (Methods). The temporal difference was computed from subtracting the previous time point intensity from the current time point intensity for each reconstructed voxel and setting the negative intensity to zero. From the temporal difference movie, we were able to observe clearly the seizure onset window, from initiation to propagation throughout the whole brain (Fig. 3d, Supplementary Video 1). The achieved imaging contrast and resolution allowed us to clearly discern the seizure activation and propagation across the various brain regions. We surmise that in future studies, the low-resolution but high volume-coverage and high-speed LFM described here could be effectively combined with other high-resolution imaging modalities, such as LSM or confocal microscopy, where LFM provides the targeted regions of interests for the latter modalities to focus the investigations on.

**Fig. 3.**
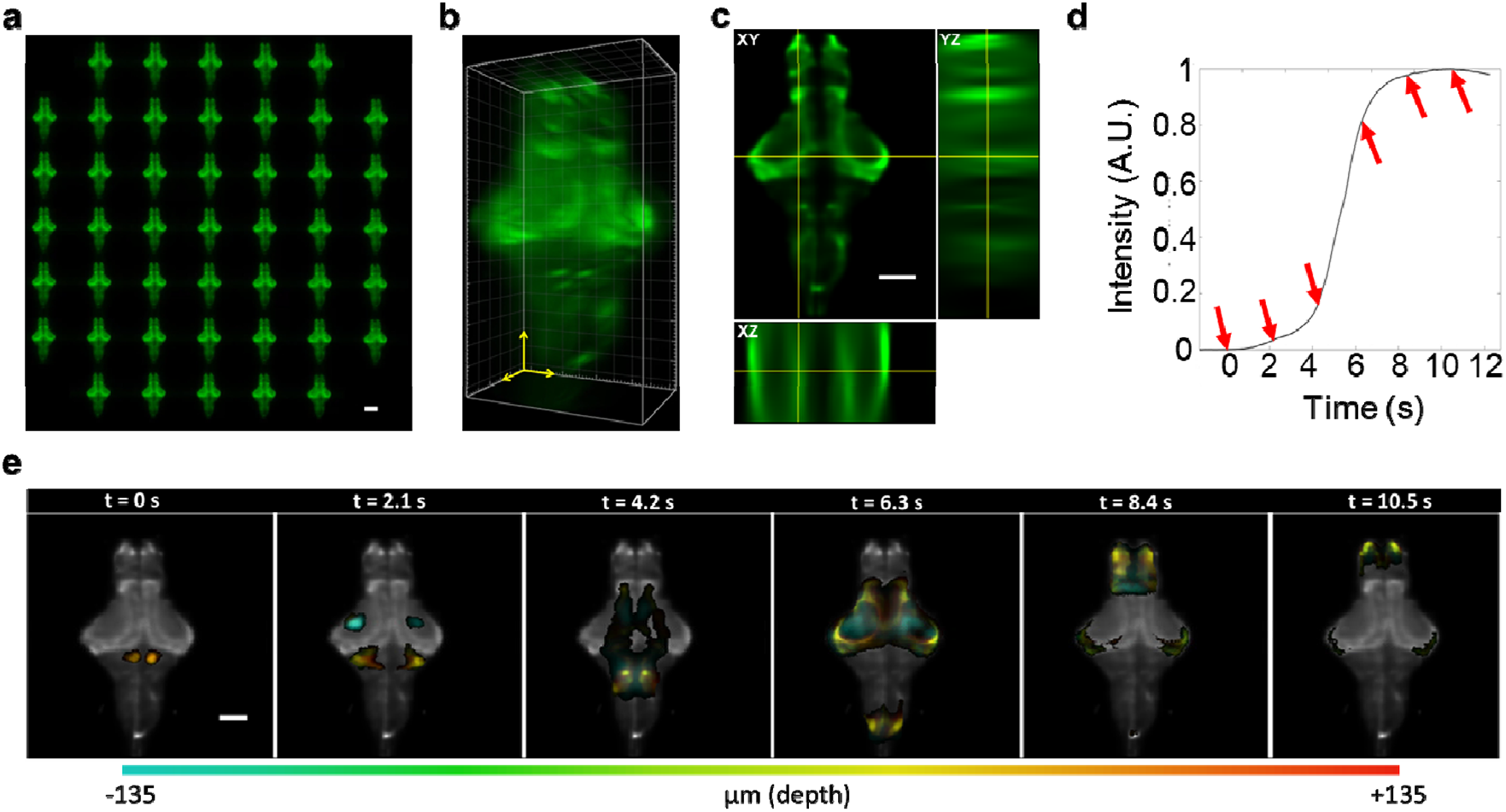
Imaging seizure in zebrafish whole brain with LFM. The fish, expressing a pan neuronal GCamp7f calcium indicator (HuC::GCaMP7f), wa s immobilized in agarose and the seizure was chemically induced by administering 15 mM PTZ (Method). The fish is at 5 days post-fertilization. The imaging is performed using 10x 0.4 NA Air immersion as the primary objective. The calcium activity was recorded at 30 volumes/s over a volume size of ∼860 × 860 × 270 µm ^3^. **a** One timeframe of FLFM recording shows different angular views of the fish brain. **b** 3D Reconstruction of a rendered in 3D. The reconstructed volume has 27 z slices with each z slice of 10 µm. The rendering shows the active neurons located in different areas of the brain. **c** Single slice cross section views of the 3D reconstructed volume in **b**. The XY view is sliced at the nominal focal plane. **d** Averaged intensity in the brain region traced over time during one seizure event. Red arrows, time points at 2.1 sec apart with the first arrow pointing at the seizure. A.U., Arbitrary Unit. **e** Depth color-coded temporal difference intensity (Method) at the time points in **c**. The seizure initialized (t = 0 sec) in the cerebellum area of the fish brain, spread out along both rostral and caudal directions. Gray scale, the average intensity over time and over axial direction presented as a contrast for the dynamical intensity. Scale bars, **a** 200 µm, **c** 100 µm, **e** 100 µm. Scale for **b** is denoted in 3 dimensions with 3 yellow arrows each of length 100 µm.

### E. Imaging calcium dynamics in mouse pancreatic islet

Calcium signaling is critical in how beta cells in pancreatic islets response to glucose, as part of their central role in maintaining glucose homeostasis and overall metabolic health^31–33^. It is generally understood that upon sensing glucose concentration above a certain threshold, beta cells in an islet spike up their intracellular calcium levels, which then kickstart a cascade leading to insulin synthesis and secretion^34–40^. Questions remain, however, on how individual beta cells coordinate their calcium activation to achieve the whole-islet response. Past imaging studies^34–40^, using conventional imaging modalities (e.g. confocal microscopy or LSM), have not resolved these questions as they lack the necessary volumetric imaging speed, topping out at ∼ 1 volumes/s to cover the entire islet^36^, while the intrinsic cellular calcium dynamic happens at sub-second time scale^41,42^. Thus, there is a need for an imaging solution that could capture calcium dynamics in the entire 3D islet at volumetric rate higher than what conventional techniques could provide, to ensure that the dynamic signals are captured faithfully.

The fast 3D imaging capability of LFM provides a solution for the above challenge. Using the 30x objective, we imaged an entire mouse islet, where the beta cells were labeled with the calcium indicator GCamp6f, in 11 mM glucose concentration, at 20 volumes/s, covering the entire volume of the islet of ∼ 100 × 100 × 86 (XYZ) µm^3^ (Methods). As shown in a representative 70-second recording (Fig. 4a, b, Supplementary Video 2), we captured the fast calcium dynamics that spontaneously initiated in a sub-region then spread in 3D across the entire islet. From the spatially-integrated intensity trace over time (Methods), we observe that the islet, as a whole, underwent repetitive calcium activation events (Fig. 4c). Within each activation event, lasting for ∼ few seconds, the fast 3D imaging speed of LFM revealed a complex spatiotemporal calcium dynamic that evolved at sub-second scale across the 3D volume of the islet. By focusing on three representative activation events and representing the calcium dynamic through the temporal difference intensity (Fig. 4d), we observe that the three calcium events initiated at different locations in the islet, then the activation spread across the islet in different 3D patterns. The observed heterogeneity in where the calcium events initiated, and how they propagated across the islet, highlights the benefits of LFM in providing the necessary fast volumetric imaging speed to capture these complex spatiotemporal dynamics. We envision our LFM platform would be the ideal imaging tool in future studies aimed at more in-depth and functional characterization of the pancreatic islet calcium dynamics, e.g. in response to changing glucose levels, and/or in concurrent recordings of insulin synthesis and release.

**Fig. 4.**
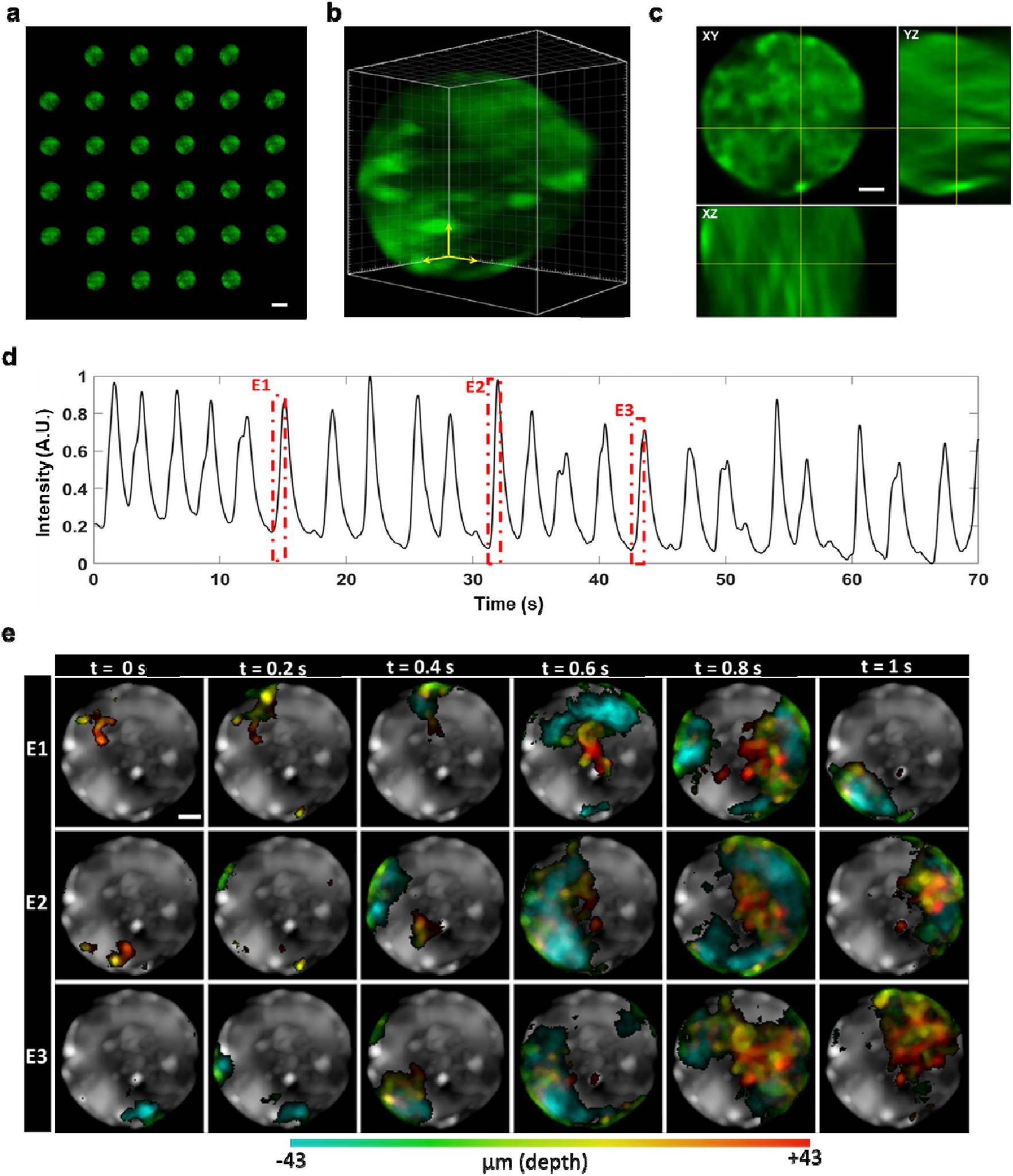
Imaging calcium dynamics in mouse pancreatic islets. Functional imaging of a mice islet with calcium indicator GCaMP6f expressed in pancreatic beta cells. The islet was in equilibrium with 11 mM glucose concentration. Imaging was performed using the 30x 1.0 NA Silicone Oil objective, recorded at 20 volumes/s. **a** Single image frame of raw LFM recording, showing the different angular views of the islet. **b** Reconstruction of **a** rendered in 3D. The reconstructed volume has 43 z-slices separated by 2 µm/slice. The unit grid is 10 µm for each dimension. **c** Single slice cross section views of the 3D reconstructed volume in **b**. The XY view is sliced at the nominal focal plane. **d** Temporal evolution of the fluorescence intensity, averaged over the entire islet. Red dashed boxes indicate the 1-second onset window of three representative calcium events (E1, E2, E3) that are analyzed further in **e**. A.U., Arbitrary Unit. **e** Depth color-coded images, of the temporal difference intensity (Methods), at six time points for calcium event E1, E2, E3 in **d**. The sequential time points are arranged horizontally, while the calcium events are separated vertically. As seen, each event initiated at a different 3D location (t = 0 sec) and propagated through the islet in different spatial patterns. The average-intensity-projection over both time and z-depth is shown as the gray-scale image in the background, for each of E1, E2, E3, to provide spatial context for the entire islet. Scale bars, **a** 100 µm, **c** 20 µm, **e** 20 µm. Scale for **b** is denoted in 3 dimensions with 3 yellow arrows each of length 20 µm.

### F. Imaging protein dynamics in live cultured cells

Toward the finer scale, we demonstrate subcellular imaging with our LFM platform, through imaging protein dynamics in live cultured cells with the 60x objective. Fig. 5 shows our results in imaging a live Human Bone Osteosarcoma Epithelial cell (U2OS line) expressing the fluorescent protein mNeonGreen fused to the human prolactin receptor (hPRLR) (Methods). The hPRLR protein plays a crucial role in various physiological processes and related diseases, including breast cancer, autoimmune disease, and metabolic disorders^43–46^. Understanding hPRLR dynamics is essential for advancing knowledge in these areas.

**Fig. 5.**
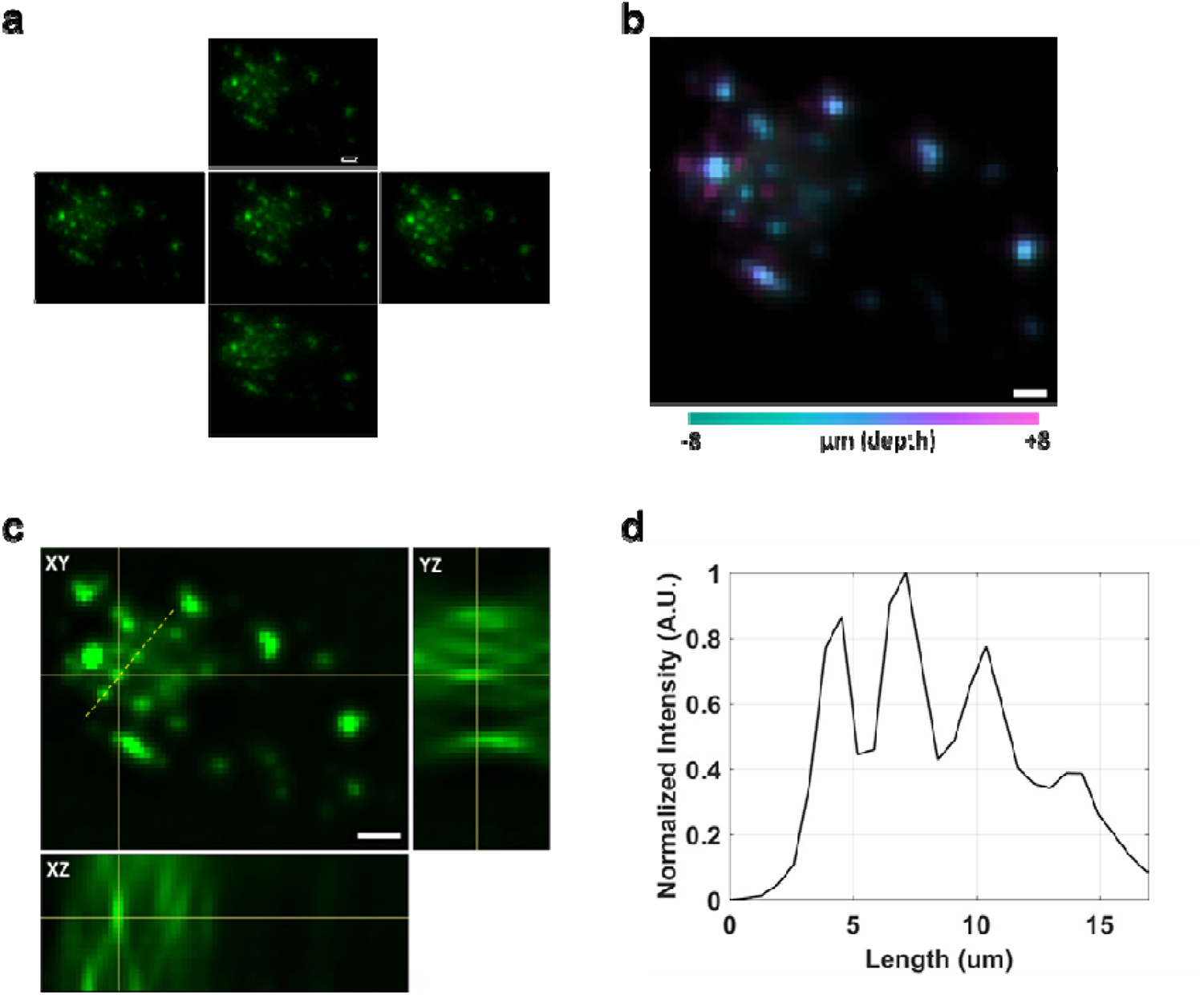
Imaging protein dynamics in live cultured cell. The Human Bone Osteosarcoma Epithelial cell (U2OS line) expressed the fluorescent protein mNeonGreen fused to the human prolactin receptor (Methods). Imaging was performed using the 60x 1.35 NA Oil objective (Table 1), recorded at 20 volumes/s (Supplementary Video 3). **a** Single image frame of the raw LFM recording, showing different angular views of the cultured cell. **b** 3D reconstruction of **a**, shown with color-coded z-depth. The reconstructed volume has 17 z-slices separated by 1 µm/slice. **c** Single slice cross section views of the 3D reconstructed volume in **b**. The XY view is sliced at the nominal focal plane. **d** Line profile of the yellow line in **c**, demonstrating the micron-level resolution in distinguishing the labeled protein puncta. A.U., Arbitrary Unit. Scale bars, 5 µm.

In the imaging experiment, we captured the rapid movement of the fluorescently-labeled hPRLR proteins throughout the cell’s volume, at volumetric speed of 20 volumes/s, over the volume of ∼ 50 × 50 × 17 (XYZ) µm^3^(Fig. 5a, b, Supplementary Video 3). Note that both the fast volumetric imaging rate of LFM and the sub-to few-micron 3D resolution achieved with the 60x objective (Table 1, Supplementary Fig. 3) were critical in this imaging scenario. High-speed or high-resolution imaging alone will not be able to faithfully resolve the dynamic movements of the subcellular protein puncta throughout the cell (Fig. 5c, d).

## III. DISCUSSION & CONCLUSION

In summary, we have presented a versatile multiscale LFM platform that is easily accessible to biological researchers. By integrating an LFM detection module with a conventional commercially available wide field microscope, we demonstrate the versatility of our LFM system, enabling observation of various 3D dynamical processes across many scales, ranging from sub-cellular protein dynamics to intercellular calcium signaling in ex vivo tissues, to whole-brain neural activity of an intact animal. The built-in flexibility and ease of use of the commercial microscope makes the LFM module accessible and user-friendly.

Unique to our LFM platform is the ability to choose between different imaging scales by simply switching to a different microscope objective, a feature shared with the previously-mentioned LFM eyepiece^23^. The two instruments, however, differ substantially in the optomechanical design and thus would excel in different application spaces. An outstanding feature of the LFM eyepiece is its portability: as the entire unit, including the recording camera, is integrated as an elongated eyepiece, it could be readily removed and applied to different microscopes, as demonstrated in^23^. In building the LFM module as an extension from the camera port of a wide field microscope (a common design choice of many commercial or lab-built add-on detection modules), our implementation trades portability for mechanical stability, where the optical components, including the camera, are bolted down with industry-standard parts. We view this choice as advantageous for applications that require high resolution and long-term stability (e.g. for timelapses), avoiding the potential stability challenges with the LFM eyepiece’s long mechanical lever arm and eyepiece-style freely-rotating press-fit mounting. Further, the open design of our LFM module has essentially no constraints on the size and weight of the camera, enabling us to use the industry-leading sCMOS camera to provide the high sensitivity needed for photon-limited bioimaging applications. Such a camera would be quite challenging to integrate with the LFM eyepiece, which is more suited for small and light cameras that could be mounted, and mechanically supported, at the end of the long lever arm. Our open design also lends itself readily to modifications and improvements, as described further below – many of these potential modifications will not be possible with the LFM eyepiece. Finally, our design leaves the standard eyepieces unchanged and available for sample view finding.

Apart from the flexible and easy-of-use design for multiscale imaging, our LFM platform is constructed as an add-on to the commercial wide microscope with all off-the-shelf optical components, making it possible for researchers without formal training in optical engineering to implement in their own lab. This concept has previously been explored by Lu et at^24^, who implemented a scanning LFM with digital adaptive optics (sLFM) to enhance imaging resolution and computationally correct for system misalignment and artifacts. While there are conceptual similarities, our work is fundamentally distinguished from their work in key hardware arrangement, and, consequently, software processing. The sLFM hardware design follows the conventional LFM architecture^9,13^, where the MLA is placed at the image plane of the optical system and the camera samples the Fourier plane. Each micro-lens image behind each micro-lens on the camera screen depicts the back aperture and NA of the objective. Under this design schematic, the NA of each micro-lens needs to be equal or slightly bigger than the NA of the objective to avoid overlapping between adjacent micro-lens images. This requirement can be met by either inserting a different relay lens for each objective or by pairing each objective with a dedicated MLA. The first option requires frequent relay-lens switching, which increases experimental complexity and user training demands, while the second often involves customized MLAs, which add cost to the implementation. Our LFM platform is designed based on the Fourier LFM (FLFM) where the MLA is positioned at the Fourier plane and the camera samples the image plane. Each micro-lens generates an angular projective view of the 3D sample with the FOV controlled by a field stop at the image plane. This effectively avoids the need for NA matching between the MLA and the objective, allowing a single MLA to be paired with multiple objectives and therefore, enabling multiscale imaging without hardware extra modifications from the user. In post processing, PSF of the FLFM is uniform across each axial plane, allowing (Fast Fourier Transform) FFT based deconvolution to be applied in the 3D reconstruction. Under this approach the reconstruction can be accelerated by approximately two orders of magnitude^14^. Compared to sLFM, our LFM system excelled in implementation simplicity, user-friendlies, and versatility, making it well-suited for bioimaging across scale.

While we built our platform on an inverted microscope body, for flexible adaptability to cells and tissues samples, the LFM module could similarly be added to an upright microscope, for applications that are more suited to that imaging configuration, such as neuroimaging. Instead of a wide field microscope, LFM can also be integrated in a similar manner with other 2D imaging systems, such as a confocal microscope or LSM, to provide an option for fast, lower-resolution synchronous volumetric imaging, to complement the existing slower, higher-resolution imaging capability of those systems. In particular, the integration of LFM with LSM enables selective volume illumination microscopy (SVIM)^47,48^, which has been shown to improve contrast and effective resolution over wide field illumination. In the work presented here, we have aimed to balance the competing needs of maximizing imaging performance versus minimizing the complexity and cost of the LFM module, thus opting for simplified optical configurations and off-the-shelf parts. If high imaging performance is the priority, different design and component choices could be made. For example, the resolution and imaging volume of the multiscale LFM platform could be enhanced with a customized MLA, with application-specific optical parameters and/or complex multi-focal features^22^. A filter-wheel-like device could be used to house different MLA’s with different specifications, thus allowing for switching in specific MLAs to match particular imaging applications. Performance enhancements could also come from a larger camera sensor size with higher pixel counts, allowing for either more angular views to be captured or higher pixel sampling rate. Further optical magnification or de-magnification could be implemented between the MLA and camera to make full use of the camera chip size, realizing the true optical resolution limit. As computing power continues to advance, our versatile multiscale LFM platform has the potential to reach more researchers and provides fast 3D imaging solutions to a broad range of biological applications, empowering scientists to explore more biological questions and unlock new working principles of living systems.

## Supporting information

Supplemental Video 1. Whole-Brain Seizure in Zebrafish Larva

Supplemental Video 2. Calcium Dynamics in Mouse Islet

Supplemental Video 3. Protein Dynamics in Cultured Cell

## ACKNOWLEDGEMENTS

We thank Dr. Amina Kinkhabwala (California Institute of Technology) and Dr. David Prober (California Institute of Technology) for protocols sharing and initial consultation for the zebrafish experiment. We thank Dr. Kate White (University of Southern California) for input to the mouse islet experiment.

## Methods

### Hardware setup and operation

The optical setup was composed of two subsystems: the commercial WFM (Olympus, IX83 Inverted Microscope) and the customized LFM module (Supplementary Fig. 1). The WFM is equipped with a fluorescence light source (Olympus, U-HGLGPS) and standard filter cube sets for different colors. The customized LFM module was appended to the left camera port of the WFM body. In the LFM module, a field stop was placed at the image plane to control the size of the field of view so that the images formed behind adjacent lenslets do not overlap each other^1–3^. The field stop limits the imaging field of view. If the user does not adjust the field stop, then adjacent micro-lens images will overlap each other. In this case, the reconstruction will not be able to reconstruct the raw FLFM image accurately. As depicted in Fig. 1a, relay lens L2, with focal length = f_2_, was positioned at distance f_2_ behind the field stop, and the MLA was placed at distance f_2_ from L2. The MLA was mounted on a customed cage plate (Thorlabs CP31with a hole machined into it to hold the MLA). To enable XY translational adjustment of the MLA, we connected the customed cage plate to a XY Translating Lens Mount (Thorlabs HPT1) through an intermediate plate (Thorlabs CP02) and a SM1 threaded adapter (Thorlabs SM1T10) (Supplementary Fig. 1). Coupled with the WFM internal tube lens L1, the relay lens L2 relayed the back aperture plane (i.e. Fourier plane) of the primary objective onto the MLA. A sCMOS camera (Hamamatsu, ORCA-Flash4.0) was placed at the back focal plane of the lenslets where the different angular views of the 3D sample were formed. For imaging in LFM mode, the fluorescence signal was directed to the left camera port in the WFM system. The emission light in all three biological samples were in the green band and thus imaged with a FITC filter cube set (Olympus, U-FF). Image acquisition was controlled by Micro-Manager^4^. A photograph of the hardware setup is shown in Supplementary Fig. 1. The key hardware components, with their specifications, are summarized in Supplementary Table 1. Add details about: (i) use objectives that have the same back focal plane position; (ii) in operation, adjust lateral position of the MLA to center the lenslets around the aperture (even vs odd # of lenslets included in the aperture).

### Zebrafish imaging experiment

*Animal model:* All zebrafish procedures were approved and completed according to the guidelines set by the University of Southern California Department of Animal Research. Zebrafish larvae in the *casper* background with pan-neuronal GCaMP7f expression, *Tg(elavl3:GCamp7f)* (gift from Dr. David Prober, California Institute of Technology), were imaged at 5 days post-fertilization. *Zebrafish preparation*: The zebrafish was paralyzed in a petri dish with 30 μL of α-bungarotoxin (MilliporeSigma, #203980) solution, at concentration of 1 mg/mL in E3 medium (5 mM NaCl, 0.17 mM KCl, 0.33 mM CaCl2, 0.33 mM MgSO4), for 30 seconds. Then 3 mL of E3 was added and the petri dish was covered in tin foil and left inside a 28°C incubator for 30 minutes. Subsequently the zebrafish was washed with 3 mL of E3 for three times, then anesthetized by placing into tricaine solution (0.05% tricaine in E3; MS 222, Sigma-Aldrich, #E10521) for a a few minutes before it was ready to be mounted for imaging. *Fish mounting:* The zebrafish was mounted in a #1.5 coverglass-bottom petri dish (MatTek, P35G-1.5-10-C), in 1.8 mL of agarose solution (1.5% agarose in E3, with 0.05% tricaine) (Thermo Fisher Scientific, UltraPure Low Melting Point Agarose, #16520100), with its dorsal side facing down and pushed as close to the bottom of the petri dish as possible. This is done to ensure the most direct optical path of the microscope objective to the brain with the inverted microscope used in the experiment. Once the agarose has solidified, the tricaine was washed out by adding and removing 3 mL of E3 for three times, with 5 minutes of equilibration time between washes. After the last wash, 1.155 mL of E3 was added. The sample petri dish was then placed in the stage-top incubator of the microscope, with the temperature set to 28°C, and left to equilibrate for at least 15 minutes. *Imaging seizures induced by pentylenetetrazole (PTZ)*: Aliquots of 45 μL (1 M) PTZ (Cayman Chemical, #18682) was made with E3 and kept frozen until use. After equilibration in the stage-top incubator, the sample was adjusted to be in the desired microscope field of view, and the focal plane was adjusted to be at ∼ 150 μm into the zebrafish brain from the dorsal surface, with microscope objective 10x 0.4 NA Air. One minute before the start of the timelapse, 45 μL (1 M) of PTZ (thawed and brought to 28°C) was added to the sample petri dish, allowing the time for closing the lid of the incubator. Timelapse was recorded for 90 minutes, with exposure time of 33.3 ms, resulting in a volumetric imaging rate of 30 volumes/s.

### Mouse islet imaging experiment

*Animal Model:* All mouse procedures were approved and completed according to the guidelines set by the Children’s Hospital of Los Angeles Institutional Animal Care and Usage Committee. Ins1Cre-GCaMP6f, β cell-specific GCaMP6f protein expression, were generated by crossing Ins1Cre mice (The Jackson Laboratory, #026801) and Ai95(RCL-GCaMP6f)-D mice (The Jackson Laboratory, #028865), both on a C57Bl/6J background. *Islet Isolation and dissociation:* Islets were isolated using a 3.5 mg/mL liberase and 1.5 mg/mL DNAse enzyme blend (Roche Diagnostics) as previously described^5^. Briefly, the enzyme solution was perfused into the pancreas via the bile duct, and the inflated pancreas was removed for further digestion at 37 °C for 12 min. Extracted islets were hand-picked under a fluorescence light microscope to isolate Ins1Cre-GCaMP6f islets and incubated with Roswell Park Memorial Institute (RPMI) 1640 (Gibco). Islets were transferred into an 8-well #1.5 coverglass-bottom imaging chamber (Ibidi, #80826) with RPMI 1640 and left at 37°C overnight before being used for experiments. On the day of imaging, islets were equilibrated in Kreb’s Ringer bicarbonate buffer as previously described^5^ at 2.8 mM basal glucose concentration, and then increased to 11 mM static glucose incubation. *Imaging*: The islet sample was placed in the microscope stage-top incubator, with temperature set to 37°C, and positioned at the center of the field of view. The focal plane was adjusted to be at the center of the islet along the z-direction, with microscope objective 30x 1.0 NA Silicone Oil. Timelapse was recorded for 70 seconds, with exposure time of 50 ms, resulting in a volumetric imaging rate of 20 volumes/s.

### Cell imaging experiment

*Plasmid design:* A plasmid containing human prolactin receptor (hPRLR), fused to the rat PRLR signal peptide, was obtained as a gift from Vincent Goffin (INSERM, Paris, France). pcDNA3 Flag HA (#10792) plasmid was obtained from Addgene. The rat PRLR signal peptide, hPRLR and the fluorescent protein mNeonGreen were cloned into the pcDNA3 plasmid using Takara In-Fusion Snap Assembly (#638947) according to the manufacturer’s protocol, such that mNeonGreen was fused to the C-terminus of hPRLR. U2OS cells were acquired from the American Type Culture Collection (Manassas, VA). *Cell culture*: Cells were cultured in DMEM (Corning, #10-013-CV), supplemented with 10% fetal bovine serum (Thermo Fisher Scientific, #A5256701), 1% penicillin/streptomycin (Thermo Fisher Scientific, #15140122), and 1% L-glutamine (Sigma-Aldrich, #G7513). For imaging experiments, cells were grown to ∼ 80% confluency before being seeded onto 8-well #1.5 coverglass-bottom imaging chamber (Ibidi, #80807). Twenty-four hours after seeding, transient transfection of the pcDNA3-hPRLR-mNeonGreen plasmid was performed using Lipofectamine 3000 and P3000 reagents (Thermo Fisher Scientific, #L30008) according to the manufacturer’s protocol. Twenty-four hours after transfection, cell culture medium was replaced with phenol red-free Leibovitz’s L-15 medium (Thermo Fisher Scientific, #21083027) for imaging. *Imaging*: The cell sample was placed in the microscope stage-top incubator, with temperature set to 37°C, and positioned at the center of the field of view. The focal plane was adjusted to be coincident with the cell, achieving the sharpest view of the sample, with microscope objective 60x 1.35 NA Oil. Timelapse was recorded for 50 seconds, with exposure time at 50 ms, resulting in a volumetric imaging rate of 20 volumes/s.

### Richardson-Lucy (RL) deconvolution

The LFM 3D reconstruction problem was solved using RL deconvolution method as described previously^6–8^. Briefly, if we arrange both the 2D image space and the 3D voxel space into vectors, the light field imaging system can be written as a matrix vector multiplication wrapped by a Poisson noise model. We denote 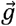 as the vectorized 3D volume, 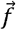 as the vectorized raw experimentally recorded 2D light field image, *Pois*{·} as the Poisson process and **H** as the PSF matrix where each column is the PSF of each voxel. The forward image formation process is characterized by the following equation:

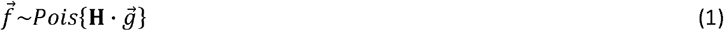

The PSF matrix **H** is acquired from the experimentally measured PSF, through imaging of sub-diffraction fluorescent beads. The goal of the reconstruction is to solve for the unknown vector 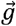. RL deconvolution was developed to solve this problem^9,10^. The deconvolution algorithm starts with an initial guess of 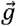, then iteratively updates the entry values in 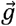 until convergence. The initiation and update are shown in equations (2) and (3):

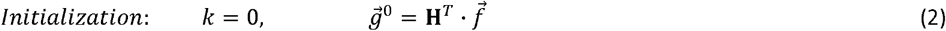

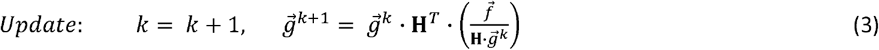

Where superscript *T* represents the transpose operator and *k* denotes the iteration times. In the update section, **H**.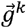 represents the forward projection process from the estimated 3D volume to the estimated 2D light field image. The pixel-wise discrepancies between the estimated 2D light field image and the true raw light field image are calculated with the term 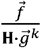, where the division denotes the element-wise division operator. The discrepancies are backward projected through 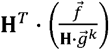 to update the estimated volume. Since the PSF is shift-invariant within each axial plane, forward and backward projection can be computed with 2D convolution using Fast Fourier Transform (FFT)^6,11^. To speed up the reconstruction, we implemented Graphics Processing Unit (GPU) acceleration. For all the reconstruction in this work, we set the total number of iterations to be 10. After 10 iterations, discernible differences were generally not observed in the reconstructed volume. Detailed discussion on the time cost of the reconstruction is provided in Supplementary Note 2.

### 3D temporal difference analysis

The temporal difference analysis is preceded by 3D spatial filtering and voxel-wise temporal filtering. First, each reconstructed 3D volume was filtered spatially by a 3D median filter of size 5 × 5 × 5 voxels. After the spatial filtering, the signal trace of each voxel was filtered by a low-pass Butterworth filter with cutoff frequency at 3 Hz. The temporal difference analysis was then applied to the filtered 3D reconstruction voxel by voxel. Let’s denote a voxel intensity value at any time point t = i as v^*i*^. Consider the timelapse that starts at t = 0 and ends at t= T. The temporal difference at t = i of each voxel is defined in (1) as:

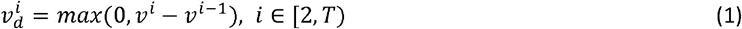

where the negative intensity was set to zero. The temporal difference was computed for all the voxels within the filtered 3D reconstructed region from the second time point to the last time point. The trace of the temporal difference represents the increase in signal intensity at each time point relative to the last time point.

