## Supplementary Notes for "Multiscale Light Field Microscopy Platform for Multi-purpose Dynamic Volumetric Bioimaging"

| Pages | Item | Description |
| --- | --- | --- |
| 2 | Supplementary Fig. 1 | Hardware set-up of the versatile LFM system |
| 3-4 | Supplementary Fig. 2 | Resolution characterization for the LFM module, with 60x and 10x primary objectives. |
| 5 | Supplementary Fig. 3 | Reconstruction speed vs linear down sampling ratio |
| 6 | Supplementary Fig. 4 | Schematic illustration of MLA illumination at Fourier plane |
| 7 | Supplementary Fig. 5 | Performance of the multiscale LFM system with two different camera pixel sizes |
| 8 | Supplementary Fig. 6 | Illustration of illuminated microlens patterns with MLA of two different MLA dimensions (N) for 60x 1.35NA Oil Objective |
| 9-12 | Supplementary Fig. 7 | Effect of the Relay and Microlens Focal Length on Performance Parameters at Different N for 30x 1.05NA Silicone Objective |
| 13 | Supplementary Video 1 | Whole-Brain Seizure in Zebrafish Larva |
| 13 | Supplementary Video 2 | Calcium Dynamics in Mouse Islet |
| 13 | Supplementary Video 3 | Protein Dynamics in Cultured Cell |
| 14 | Supplementary Table 1 | Part numbers and descriptions of key components of the LFM module |
| 15 | Supplementary Table 2 | Theoretical estimates of imaging performance |
| 16 | Supplementary Table 3 | Imaging and reconstruction parameters of presented results |
| 17 | Supplementary Table 4 | Resolution for Wide Field Microscope using different objective |
| 18-19 | Supplementary Tables 5-10 | Estimated theoretical imaging performance of the multiscale LFM system with MLAs of different number of microlenses and different camera pixel sizes |
| 20 | Supplementary Table 11 | The Number of Reconstructed Voxels vs the Number of Recorded Pixels |
| 21 | Supplementary Note 1 | Calculation of performance parameters |
| 22 | Supplementary Note 2 | Determination of number of microlenses in the MLA |
| 23-24 | Supplementary Note 3 | Experimental PSF processing and 3D reconstruction of LFM data. |

**Supplementary Figure 1. Hardware set-up of the versatile LFM system**

Light Field Microscopy (LFM) is implemented as an add-on module (yellow box) appended to a commercial Wide Field Microscope (WFM) body (cyan box). The red arrow is pointed to the objective turret integrated within the WFM. The nosepiece can house six objectives in total and is controlled automatically by the microscope controller. The field stop is mounted at the image plane of the standard camera port to control the field of view. In the LFM module, there are three main components: a relay lens (L2), a micro lenslet array (MLA), and a recording camera. The relay lens L2 has focal length f_2_ = 180 mm, same as the tube lens in the WFM. The MLA is square shaped with each side of length 12 mm. Each microlens is also square shaped with focal length f_ML_ = 29.9 mm and diameter of 1.5 mm. The MLA was mounted on a customed cage plate (Thorlabs CP31 with an aperture machined into it to hold the MLA). To enable XY translational adjustment of the MLA, we connected the customed cage plate to a XY Translating Lens Mount (Thorlabs HPT1) through an intermediate plate (Thorlabs CP02) and a SM1 threaded adapter (Thorlabs SM1T10) (green dotted box). The camera is a scientific Complementary Metal–Oxide–Semiconductor (sCMOS) camera from Hamamatsu (ORCA-Flash4.0). The chip size is 13 × 13 mm^2^ with pixel size of 6.5 µm, rendering an image size of 2048 × 2048 pixels. A list of all the hardware models and parameters are listed in Supplementary Table 1.

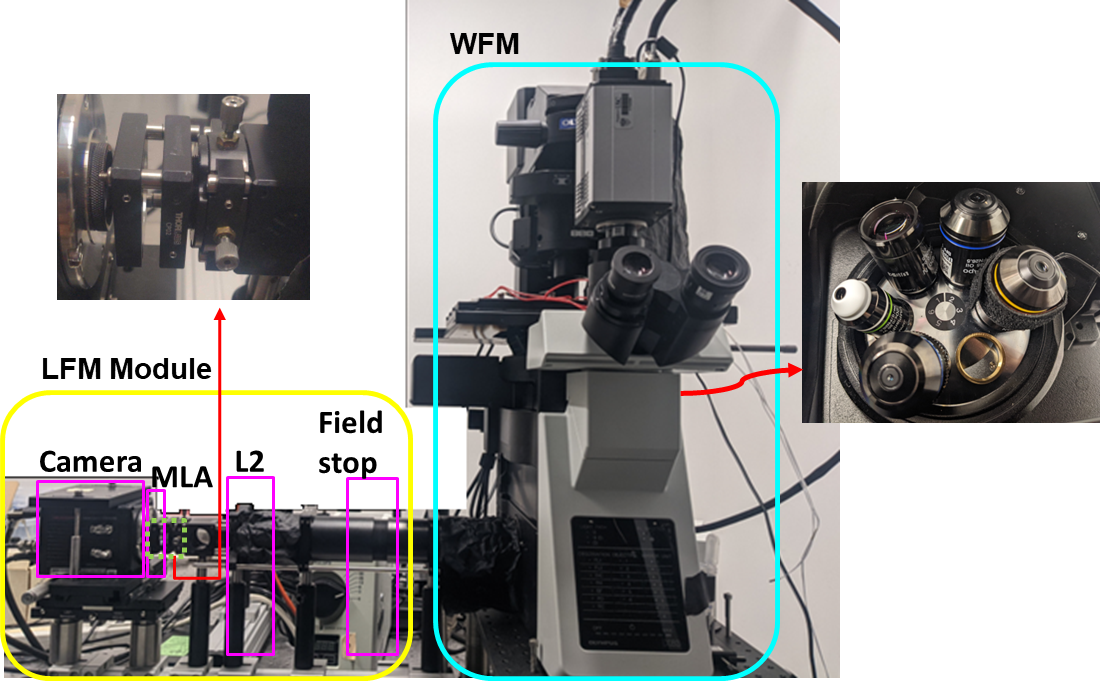

**Supplementary Figure 2.** **Resolution characterization for the LFM module, with 60x and 10x primary objectives.**

**a & b** Maximum-Intensity Projection of a 1-µm-diameter fluorescent bead image z stack, taken with **a** 60x 1.35 NA Oil objective and **b** 10x 0.4 NA Air objective, color-coded for z-depth over the depth-of-view from -8 µm to 8 µm and -135 µm to 135 µm, respectively. The angular view behind each microlens is separated into yellow circles. Four zoomed-in views of white boxes 1-4 are shown on the right side.

**c & d** Lateral and axial profiles of the reconstructed bead from **a & b,** respectively. A.U., Arbitrary Unit.

**e** FWHM measurements compared to theoretical estimates of the resolutions, for the 60x 1.35 NA Oil, 10x 0.4 NA Air and 30x 1.05NA Silicone objective. FWHM of five reconstructed beads were measured and plotted individually as asterisks (*). The FWHM averages are shown as solid dots, and the error bars show the standard deviations. Theoretical estimates of the lateral and axial resolution are presented as cyan and magenta columns, respectively (Table 1). Overall, experimental resolution values agreed, and exceeded, theoretical estimates (see text for details). Scale bars, (**a, c**) 5 µm, (**b**) 100 µm, (**d**) 10 µm.

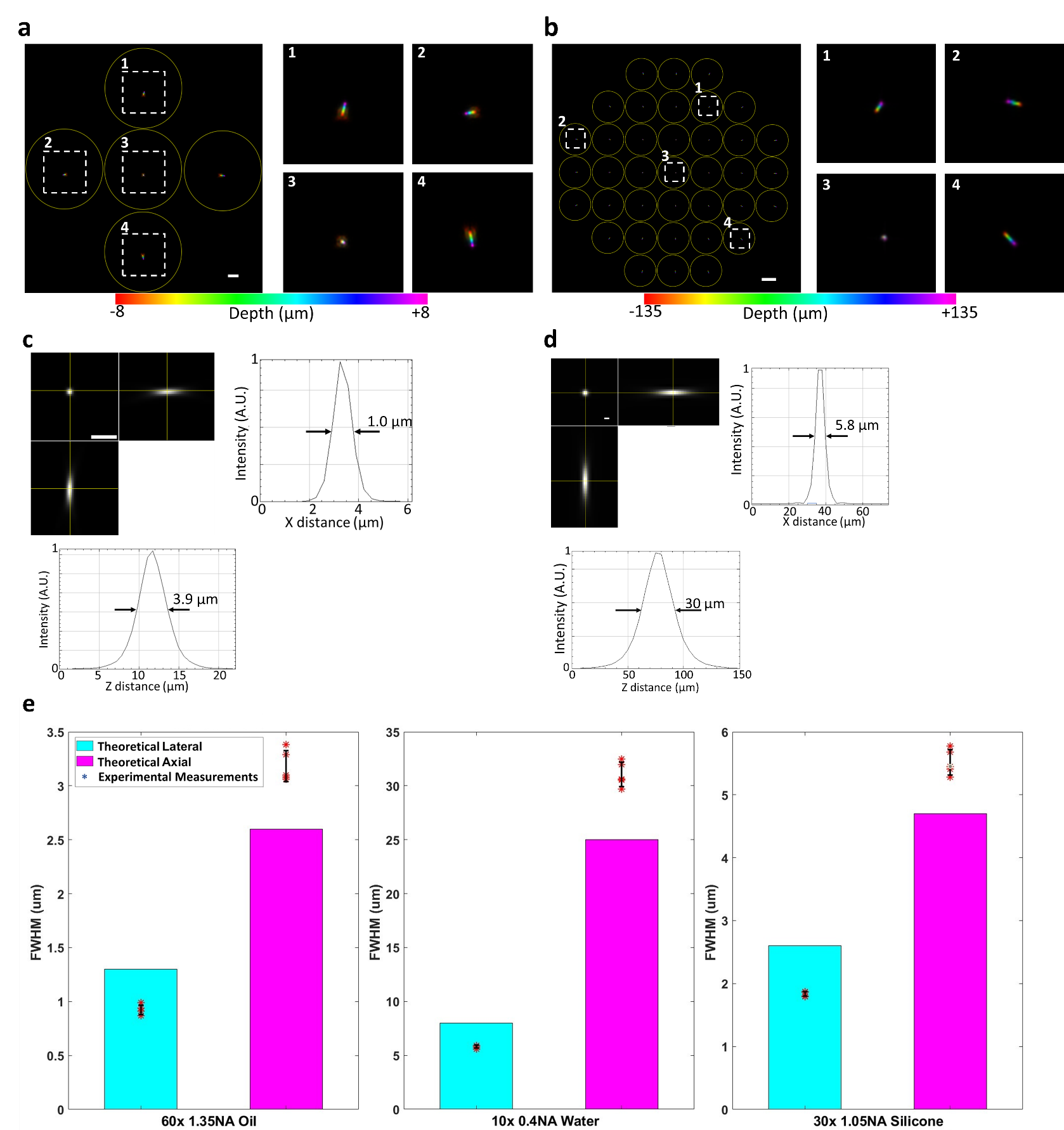

**Supplementary Figure 3. Reconstruction speed vs linear down sampling ratio**

The reconstruction time is analyzed with respect to the size of a raw 2D light field image with 61 reconstructed z-slices. The linear down sampling ratio (LDSr) is defined as the down-sampling ratio for each side of an image. For a light field image of initial size 2048 × 2048 pixels, the down-sampled image would have size: (2048 × LDSr) × (2048 × LDSr) = LDSr^2^ × 2048 × 2048 pixels. The number of the reconstructed z-slices would be *round* (61 × LDSr), where *round* (·) represents rounding to the nearest integer.

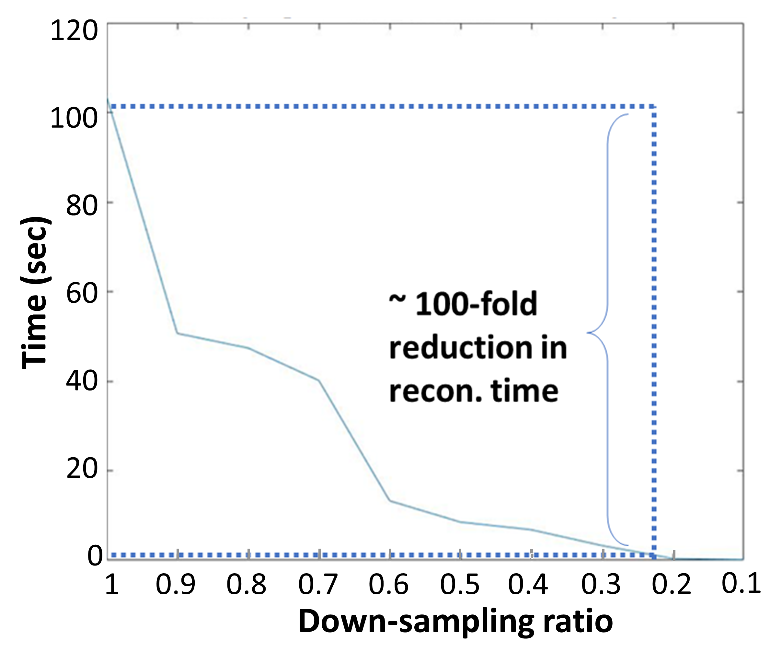
The reconstruction is performed at 5 iterations per image with the GPU Quadro P5000. As highlighted in the dotted blue lines in the plot, to have ~ 100-fold reduction in reconstruction time, the LDSr would be ~ 0.22. Without down-sampling, a 5-min recording, taken at 100 frames per second, requires 84 hours (3.5 days) to reconstruct. After the down-sampling with LDSr = 0.22, the reconstruction takes less than an hour.

**Supplementary Figure 4. Schematic illustration of MLA illumination at Fourier plane.**

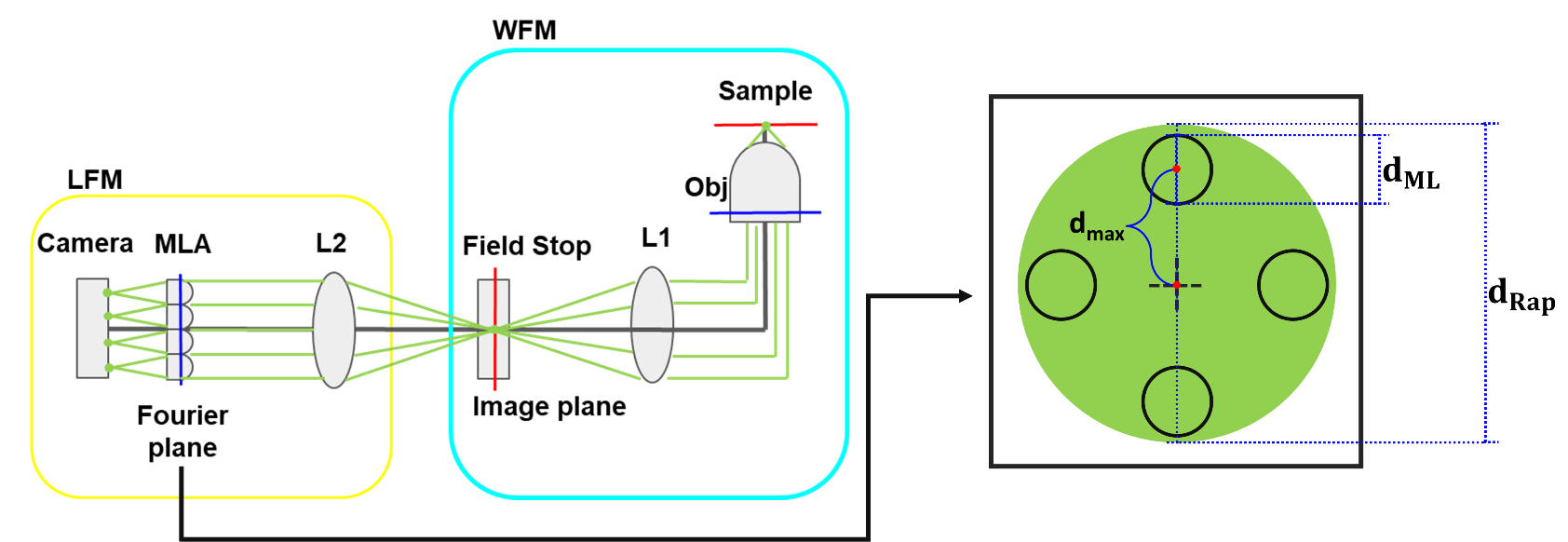
The back aperture of the objective is relayed onto the MLA, which gives us access to the Fourier plane of the optical system. Since the focal lens of L2 and L1 are the same, this relay is a one-to-one relay without magnification factor. Denote the Relayed aperture size as $ⅆ_{\mathrm{Rap}}$. $ⅆ_{\mathrm{Rap}}$ is computed as $ⅆ_{\mathrm{Rap}}= \frac{2NA\cdot f_{2}}{M}$. The effective NA at each micro-lens is reduced by a factor of $N_{r}=\frac{ⅆ_{\mathrm{Rap}}}{ⅆ_{\mathrm{ML}}}=\frac{2NA\cdot f_{2}}{M\cdotⅆ_{\mathrm{ML}}}$. The distance between the aperture center and the outmost micro-lens is denoted as $ⅆ_{\max}$, which is an important hardware parameter for computing the resolution as shown in Supplementary Table 2.

**Supplementary Figure 5. Performance of the multiscale LFM system with two different camera pixel sizes.**

Performance of all microscope objectives of interests, represented through the single resolvable voxel size and number of resolvable voxels, as a function of N. Note that here, for each objective, N starts at the minimum MLA dimension (equivalently, the maximum microlens pitch $d_{ML}=\frac{Diameter of MLA}{N}=\frac{12}{N}$) where at least at least 2 × 2 number of microlenses are illuminated. Taking the 60X 1.35 NA Oil objective as an example: at N = 6 ($d_{ML}=\frac{12}{6}=2 mm$), as shown in Supplementary Fig 6a, the 5.3 mm diameter of the back aperture allows a 2 × 2 array of microlenses to be fully illuminated. For any N < 6 ($d_{ML}>2 mm$ a single 1 × 1 microlens can be fully illuminated, which eliminates the light field effect. The performance parameters were calculated using equations (1) and (2) in Supplementary Note 1. The objectives vary over a wide range, from 10X 0.4 NA Air to 60X 1.35 NA Oil (Table 1). The averages over all the objectives’ performance curves are plotted in black. **a, c** The performance for camera pixel size of 2.2 µm. As N decreases, the number of Resolvable Voxels increases, and the voxel size decreases monotonically. **b, d** The performance for camera pixel size of 6.5 µm. As N decreases, the number of voxels increases and the voxel size decreases, but both reach a plateau around N = 10. The performance curves for the plateau region N = ~ 6-10, is a direct effect of the larger pixel size (6.5 µm), which dominates the resolution limit due to the Nyquist sampling rate, despite continued improvements in optical resolution. In our implementation, we used a camera with a 6.5 µm pixel size and an MLA dimension at N =8. While this configuration is not the absolute optimum for performance, it was chosen based on the commercial availability as well as what’s commonly used in many labs. In the optimal case, a smaller pixel size, closer to 2.2 µm, together with a smaller N value should be used to achieve the optimal optical performance limit.

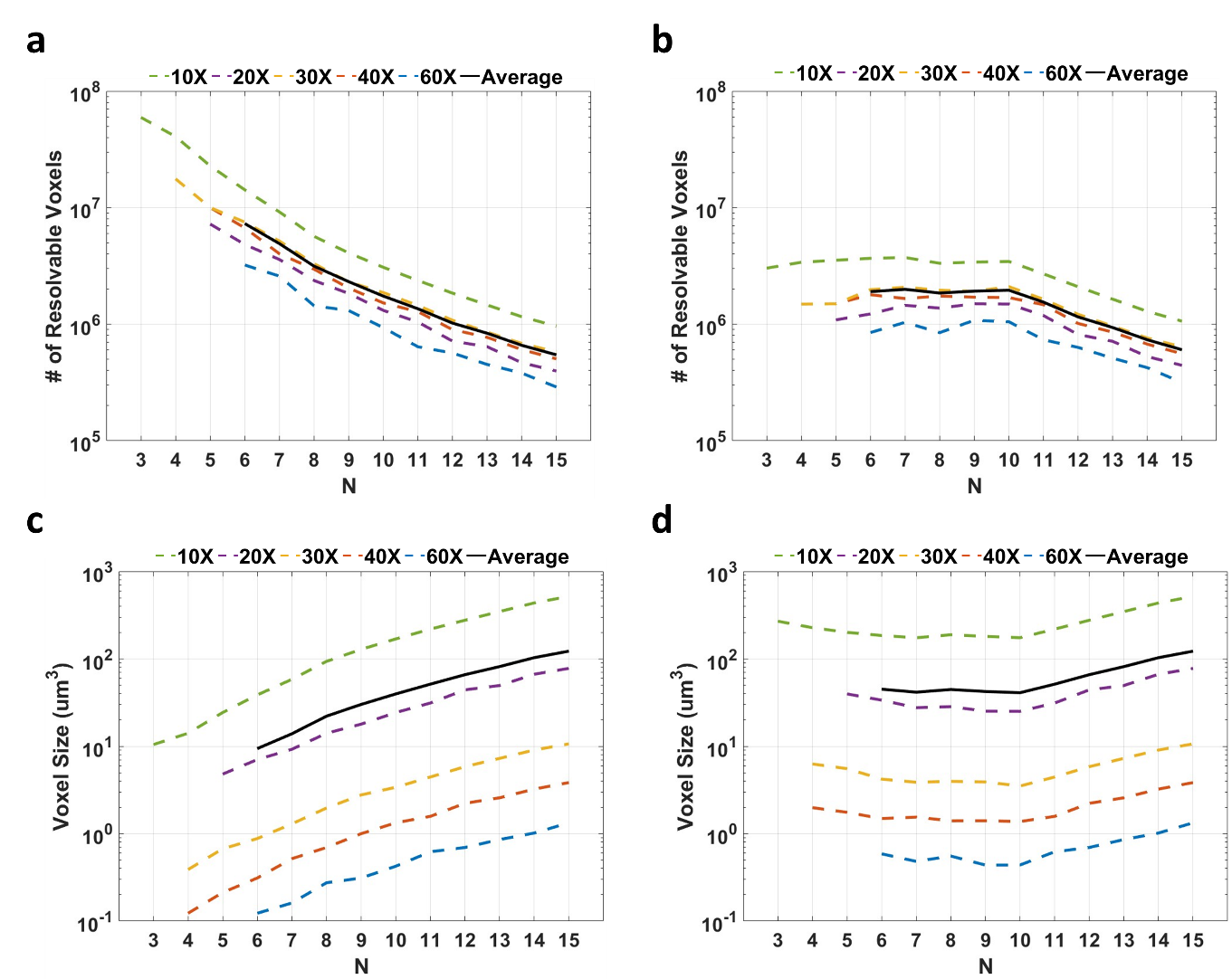

**Supplementary Figure 6. Illustration of illuminated microlens patterns with MLA of two different MLA dimensions (N) for 60x 1.35NA Oil Objective.**

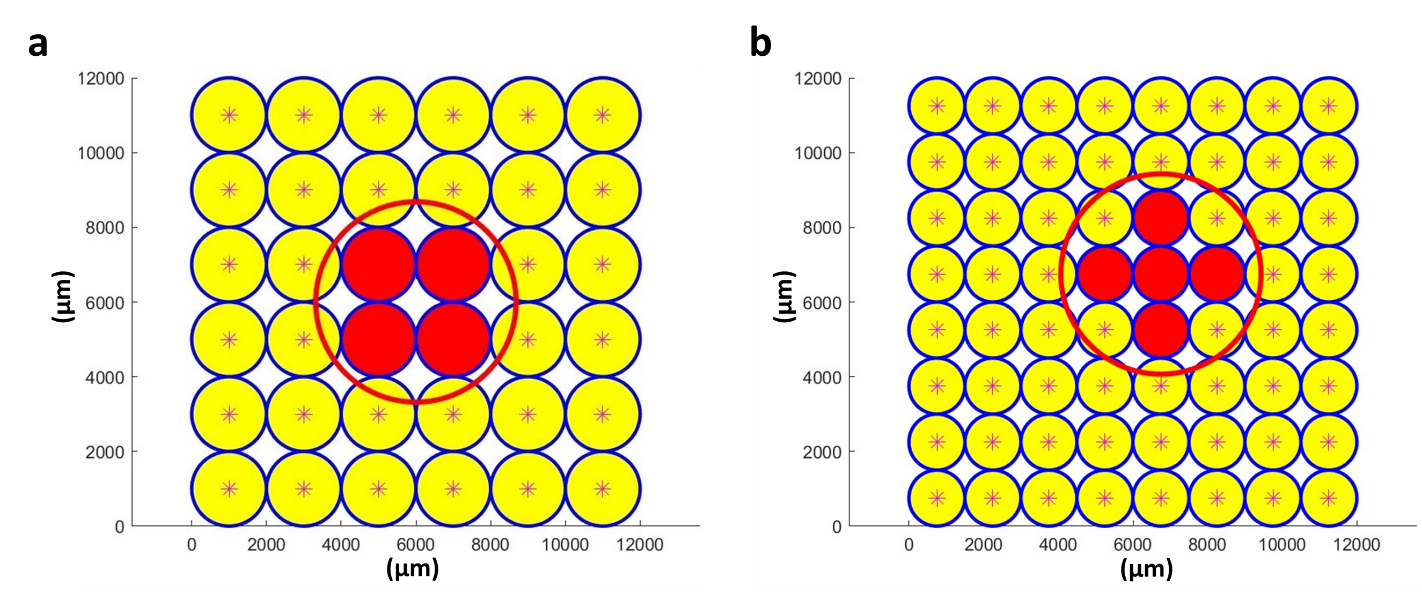
The back aperture of the objective is relayed onto the MLA. In both system design and 3D reconstruction, only fully illuminated microlens images within the relayed back aperture are considered. The outline of the relayed back aperture light is depicted in a red circle, and the center of the aperture is marked in a large blue star. The size of the MLA is 12 × 12 µm. Each microlens is represented as a yellow circle with blue outline. The fully illuminated ones are painted in red circles with blue outlines. The center of each microlens are marked in red star (*). **a** MLA with microlens dimension of 6 × 6 (N = 6). **b** MLA with microlens dimension of 8 × 8 (N = 8). Comparing **a** & **b**, N results in different microlens diameter, and therefore different number of microlens fully illuminated. For smaller N, the diameter of each microlens is bigger, leading to fewer microlens images being fully illuminated (2 × 2 for N = 6 versus 3 × 3 – 4 for N = 8). In this work, we consider the light field effect to be meaningful when there are at least a 2 × 2 array of microlenses to be fully illuminated. This determines the starting point of the performance plots with respect to N for each objective shown in Fig 1b & c as well as in Supplementary Fig 5.

**Supplementary Figure 7. Effect of the Relay and Microlens Focal Length on Performance Parameters at Different N for 30x 1.05NA Silicone Objective**.

f_2_ denotes focal length of the relay lens L_2_ (Supplementary Fig 1). f__ml_ denotes focal length of each microlens. **a-l** Color-coded performance plots for total number of resolvable voxels and single voxel size with respect to f_2_ and f__ml_ for N from 4 to 15. For each N, total number of resolvable voxels is plotted on the left and the single voxel size is on the right. Since the size of the objective back aperture is relayed and magnified through relay lens L_2_ onto the MLA at a magnification factor of $\frac{f_{2}}{f_{1}}$, each plot starts at the minimum $f_{2}$ where at least 2 × 2 number of microlenses are fully illuminated (Supplementary Fig 6a). Here, $f_{1}$ is the tube lens for the objective and is fixed for each objective. For our setup at N = 8 in **e**, the optimal f_2_ and f__ml_ for the total number of resolvable voxels and the single voxel size are marked in red and cyan crosses respectively, which represents the set of f_2_ and f__ml_ that yields the greatest number of resolvable voxels and the least single voxel size. Ideally, a good choice of f_2_ and f__ml_ would be in the middle area of the two crosses. Our choice of f_2_ and f__ml_ are marked in white cross. It is not the exactly optimal for 30x objective. However, our main goal is to try to find a balance for all the objectives as well as providing a performance metric and evaluation pipeline for exploring the potential system design.

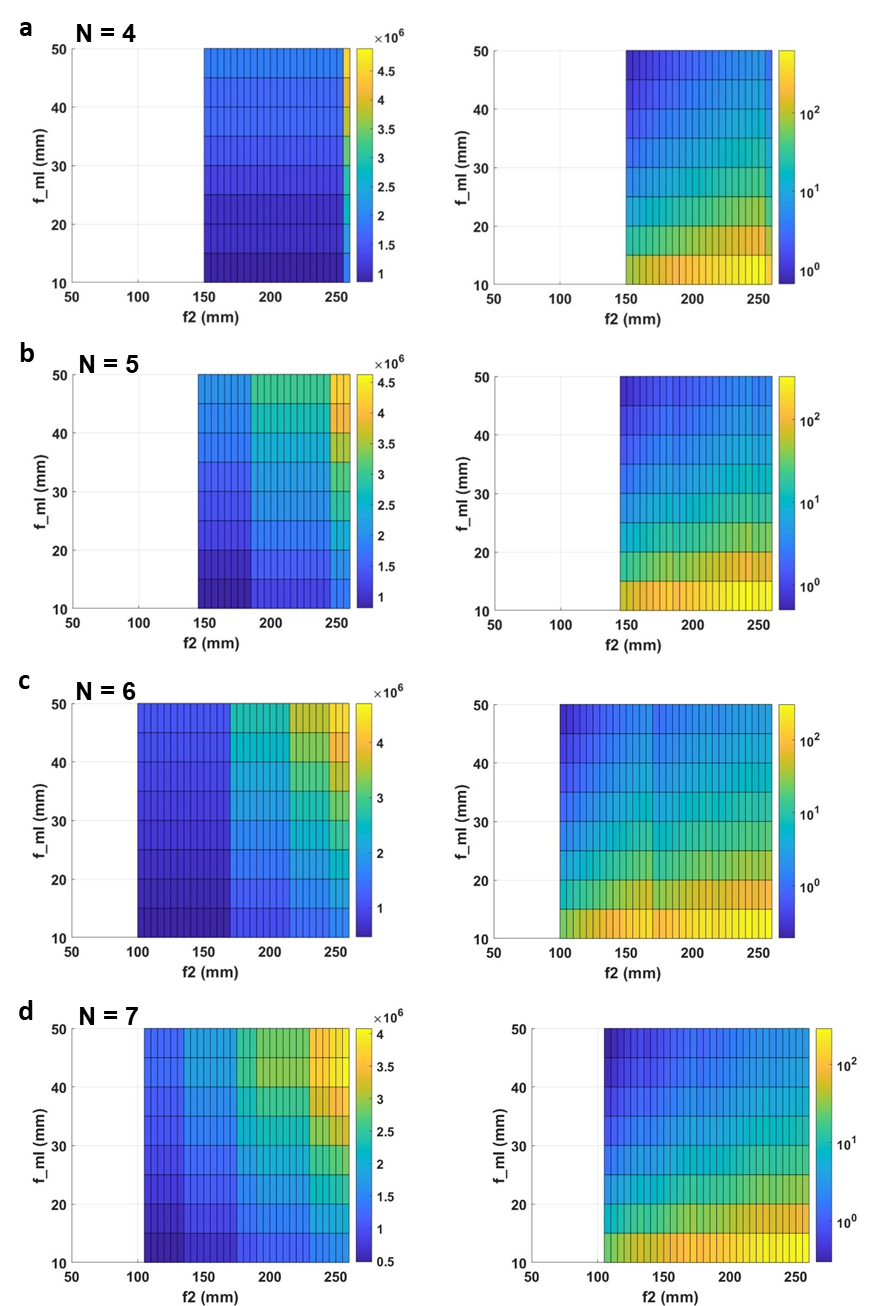

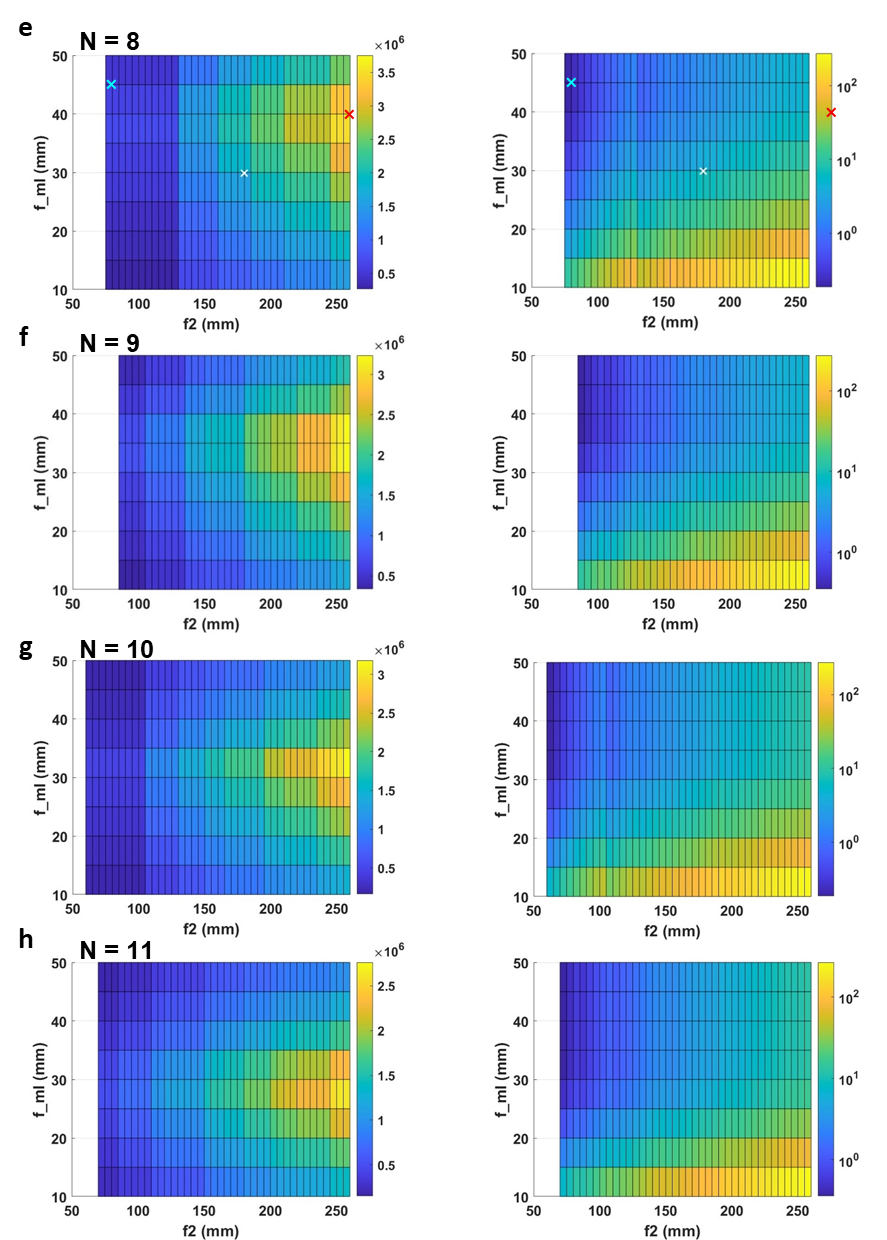

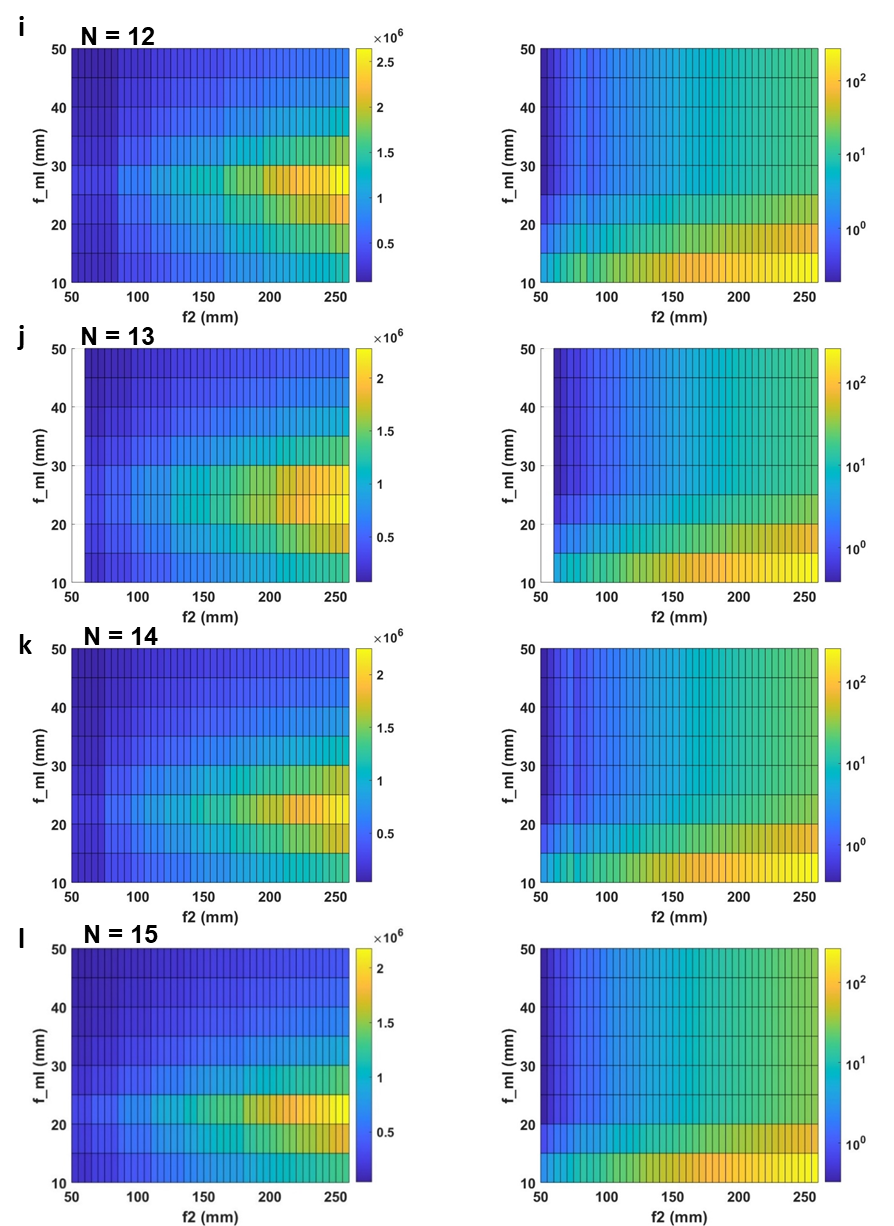

**Supplementary Video 1. Whole-Brain Seizure in Zebrafish Larva**

Whole-brain LFM imaging of a 5-dpf zebrafish larva expressing a pan-neuronal calcium indicator, *Tg(elavl3:GCaMP7f).* The sample was immobilized in agarose and the seizure was chemically induced by administering 15 mM PTZ. The seizure event was recorded at 33 volumes/s.

**Supplementary Video 2. Calcium Dynamics in Mouse Islet**

LFM imaging of an ex vivo pancreatic islet, from a transgenic mouse where all the pancreatic beta cells express the calcium indicator GCaMP6f. The islet was in a medium with glucose concentration at 11 mM. The calcium dynamic activity was recorded at 20 volumes/s.

**Supplementary Video 3. Protein Dynamics in Cultured Cell**

LFM imaging of the 3D protein dynamics in a cultured U2OS cell expressing mNeonGreen fused to the human prolactin receptor. The dynamics were recorded at 20 volumes/s.

**Supplementary Table 1. Part numbers and descriptions of key components of the LFM module**

| **Item** | **Acronym**  **(Sup. Fig. 1)** | **Descriptions** | **Part number** | **Manufacturer/**  **Vendor** |
| --- | --- | --- | --- | --- |
| Wide Field Microscope | WFM | Inverted WFM with motorized stage  Light source: Olympus, U-HGLGPS  Filter cube set: Olympus, U-FF | IX-83 Inverted | Olympus |
| Field stop | Field stop | Ring-Actuated Iris Diaphragm  Aperture range: Ø1.4 - Ø25.0 mm | SM2D25D | Thorlabs |
| Relay lens | L2 | Focal length = 180 mm | AC508-180-A | Thorlabs |
| Micro-lens array | MLA | Pitch = 1.5 mm,  Focal length = 29.9 mm,  Area = 12 × 12 mm^2^ | APD-GB-P1500-R13.8 | OKO Tech |
| LFM Camera | Camera | Scientific CMOS  Pixel size = 6.5 µm, 2048 × 2048 pixels | ORCA-Flash4.0 | Hamamatsu |

**Supplementary Table 2. Theoretical estimates of imaging performance.**

| Performance Parameters | Governing Equations | Hardware Parameters* |
| --- | --- | --- |
| FOV | $\frac{ⅆ_{\mathrm{ML}}\cdot f_{2}}{f_{\mathrm{ML}}\cdot M}$ | $\lambda$ : Wavelength of fluorescence  $p$ : Linear size of camera pixel  $ⅆ_{\mathrm{ML}}$ : Diameter of each microlens of the MLA  $ⅆ_{max}$: Distance from the aperture center to the microlens with maximum angular view (outermost microlens) **  $f_{\mathrm{ML}}$ : Focal length of each microlens  $f_{2}$ : Focal length of Relay lens L_2_  $M$ : Magnification of objective  n : reflective index of objective immersion media |
| DOV | $\frac{8\lambda\cdot{f_{2}}^{2}}{M^{2}{{\cdotⅆ}_{\mathrm{ML}}}^{2}}+\frac{4p\cdot{f_{2}}^{2}}{M^{2}\cdotⅆ_{\mathrm{ML}}{\cdot f}_{\mathrm{ML}}}$ |  |
| $R_{xy}$ | $max(\frac{2\lambda\cdot f_{2}}{n{\cdot M\cdotⅆ}_{\mathrm{ML}}} , 2p\cdot\frac{f_{2}}{{M\cdot f}_{\mathrm{ML}}})$ |  |
| $R_{z}$ | $R_{xy}\cdot\frac{f_{2}}{{M\cdot d}_{max}}$ |  |

* All samples emit fluorescent light at $\lambda=0.516 nm$. $M$ and n are determined by specific objectives. Values of the hardware parameters in the light field module are: $p=6.5 \mu m$; $ⅆ_{\mathrm{ML}}=1.5 \mathrm{mm}$; $f_{\mathrm{ML}}=29.9 \mathrm{mm}$; $f_{2}=180 \mathrm{mm}$.

$ⅆ_{max}$ varies with the back aperture size of each objective. The values are listed below:

| Objective | 60x, 1.35 NA | 40x, 1.25 NA | 30x, 1.05 NA | 20x, 0.5 NA | 10x, 0.4 NA |
| --- | --- | --- | --- | --- | --- |
| $ⅆ_{max}$ (mm) | 1.5 | 3.0 | 3.4 | 2.4 | 5.7 |

**The illustration of $ⅆ_{max}$ is shown in Supplementary Fig. 4.

**Supplementary Table 3. Imaging and reconstruction parameters of presented result**

|  | LFM image size | Reconstructed number of Z slice | Number of frames | Acquisition time per frame  (milliseconds) | Reconstruction time per frame (seconds) | Total Reconstruction time  (hours) | Total acquisition time  (seconds) |
| --- | --- | --- | --- | --- | --- | --- | --- |
| Zebrafish brain  (Fig. 3) | 1775 × 1775 | 27 | 451 | 33.3 | 20 | 2.5 | 15 |
| Mice islet  (Fig. 4) | 1395 × 1395 | 43 | 1400 | 50 | 23 | 9 | 70 |
| Cultured cell  (Fig. 5) | 713 ×  713 | 21 | 1000 | 50 | 1.7 | 0.5 | 50 |

**Supplementary Table 4. Resolution for Wide Field Microscope using different objective**

| Objectives | Magnification | 60 | 40 | 30 | 20 | 10 |
| --- | --- | --- | --- | --- | --- | --- |
|  | Numerical Aperture | 1.35 | 1.25 | 1.05 | 0.5 | 0.4 |
|  | Refractive Index | 1.51 | 1.41 | 1.41 | 1.33 | 1.00 |
|  | Wide Field Microscope Resolution (µm) | 0.11 | 0.16 | 0.22 | 0.33 | 0.65 |

**Supplementary Tables 5-10. Estimated theoretical imaging performance of the multiscale LFM system with MLAs of different number of microlenses and different camera pixel sizes.**

PX: pixel sampling resolution. DL: diffraction limited optical resolution. When pixel sampling rate is below 2, which is the Nyquist sampling rate, the resolution is determined by the pixel sampling resolution. When pixel sampling rate exceeds 2, the system resolution is capped by diffraction limited resolution. In each Table, the final system resolution is highlighted in green. Comparing Table 5 and 7, we could see that pixel resolution is determined by the pixel size independent of the number of microlens in the MLA (equivalently the microlens pitch). Comparing Table 5&6 and 7&8, we could observe that diffraction limited resolution is determined by the number of microlenses in the MLA independent of the pixel size.

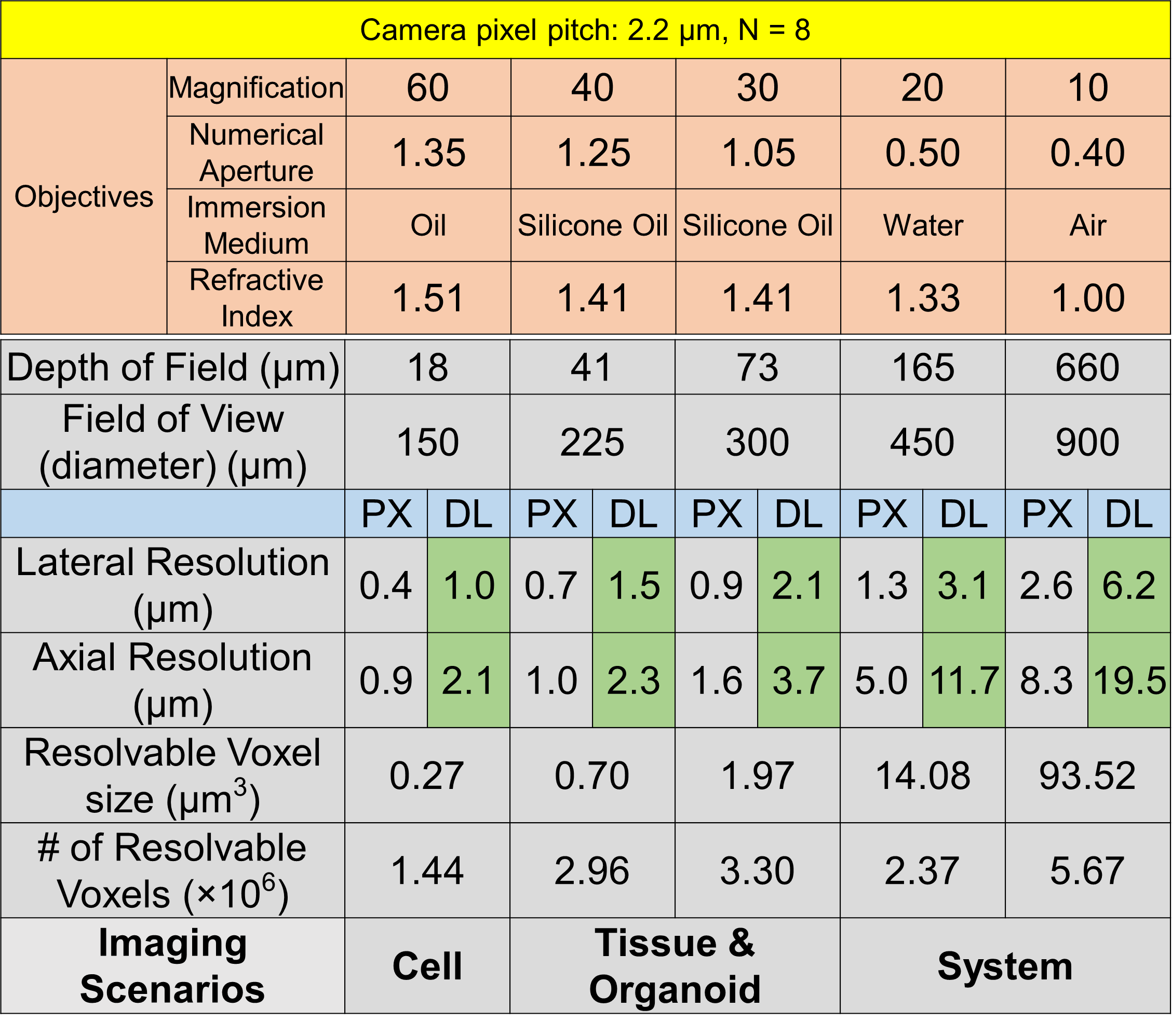

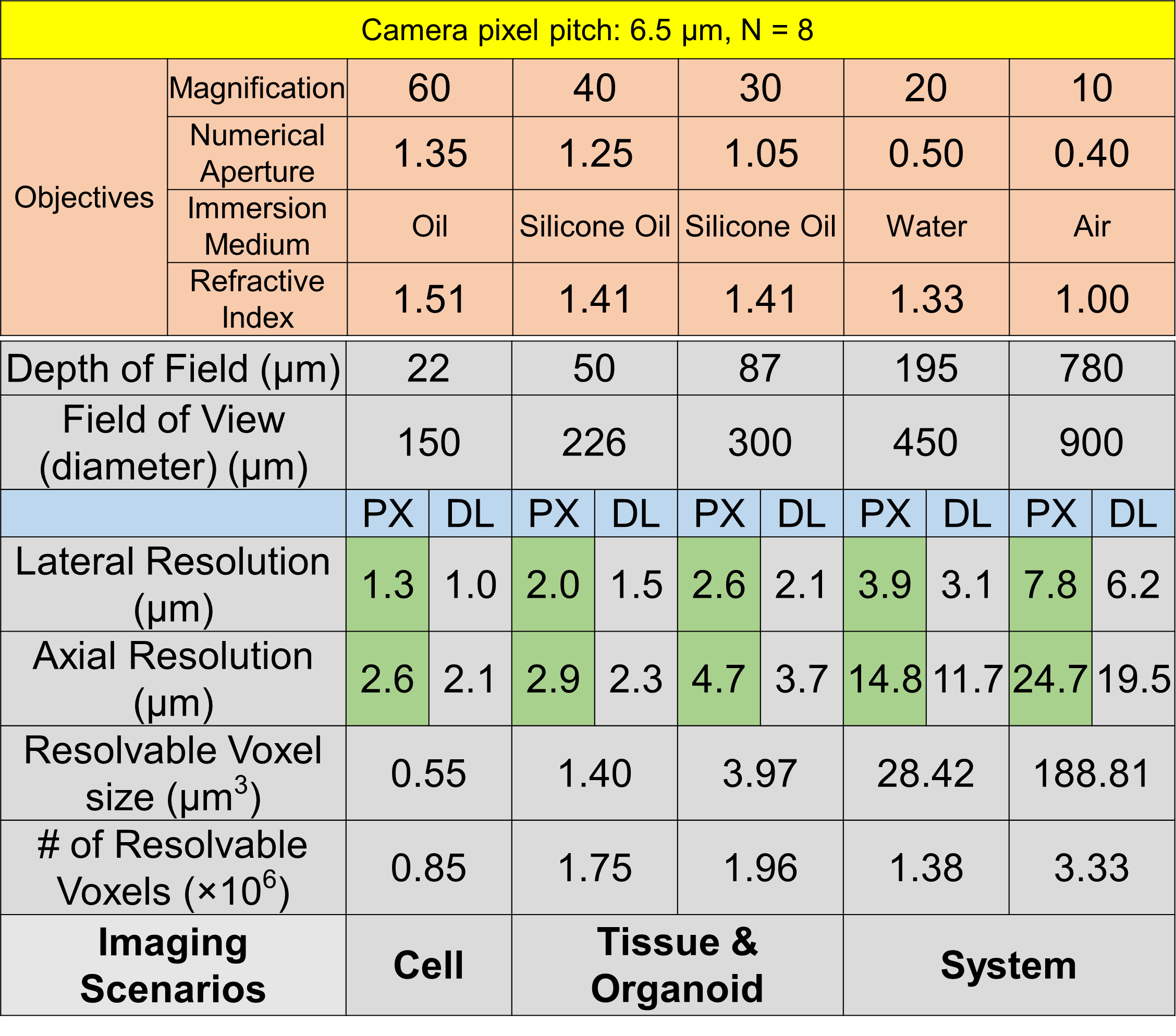

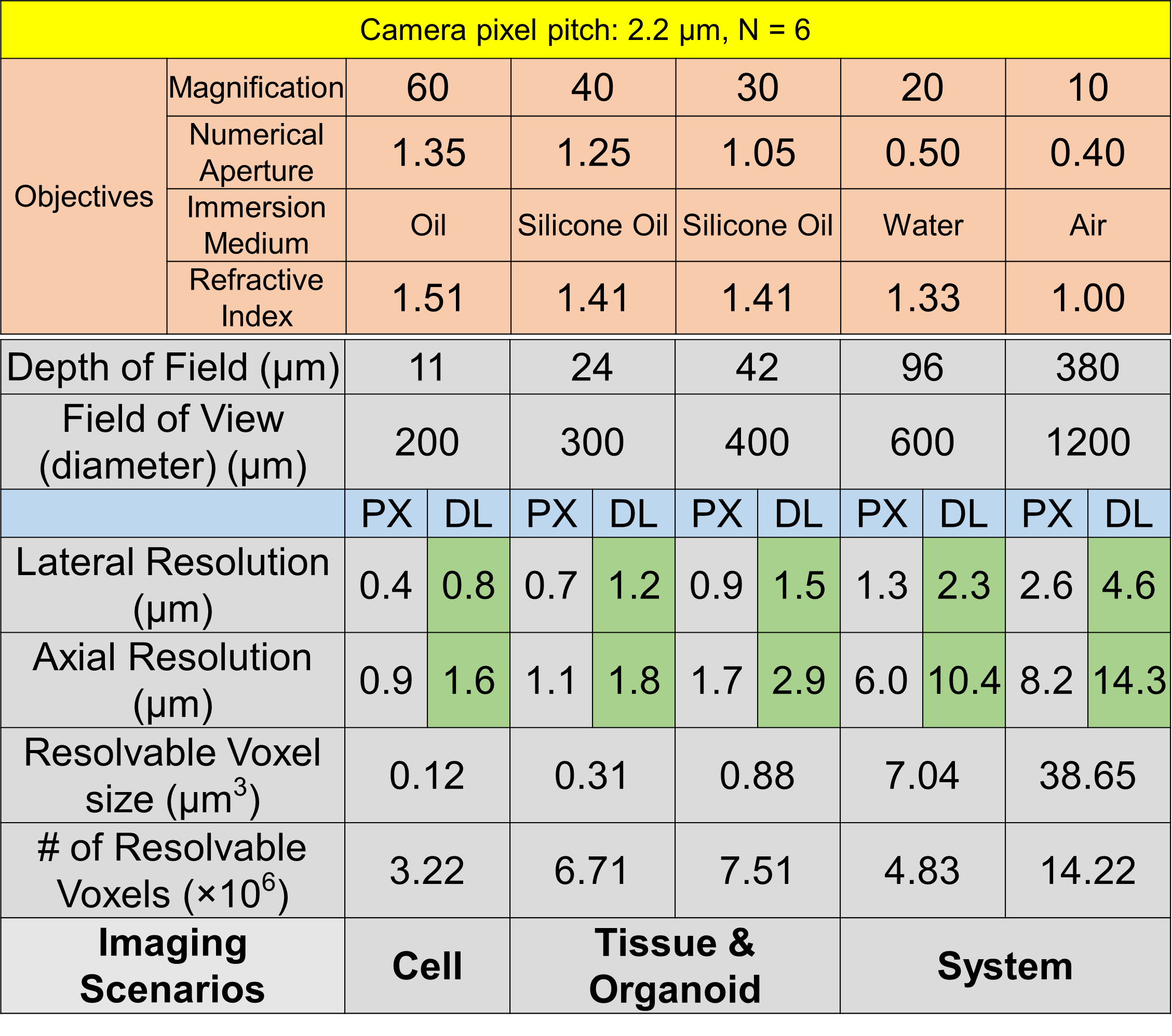

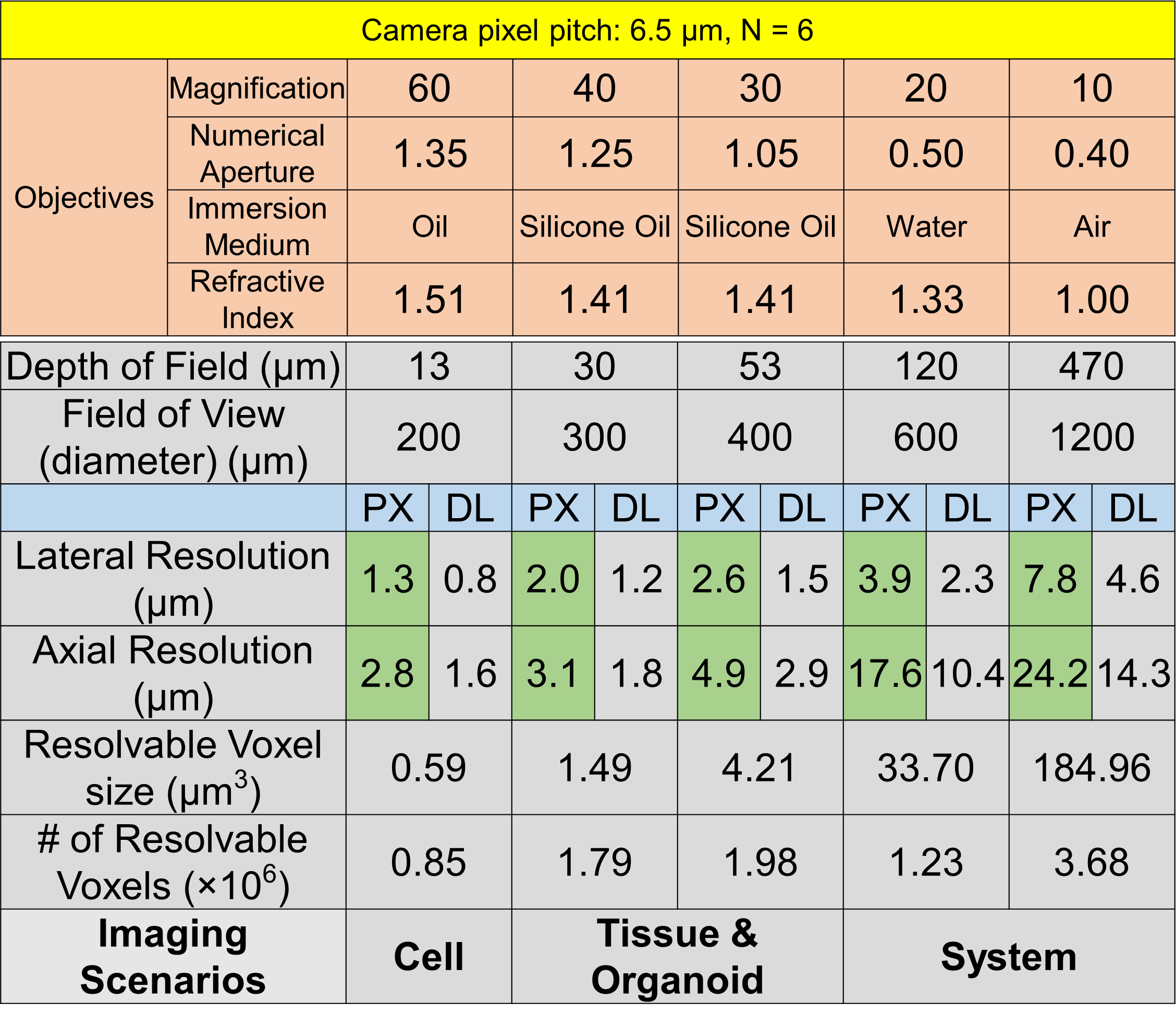

**Table 5**

**Table 6**

**Table 8**

**Table 7**

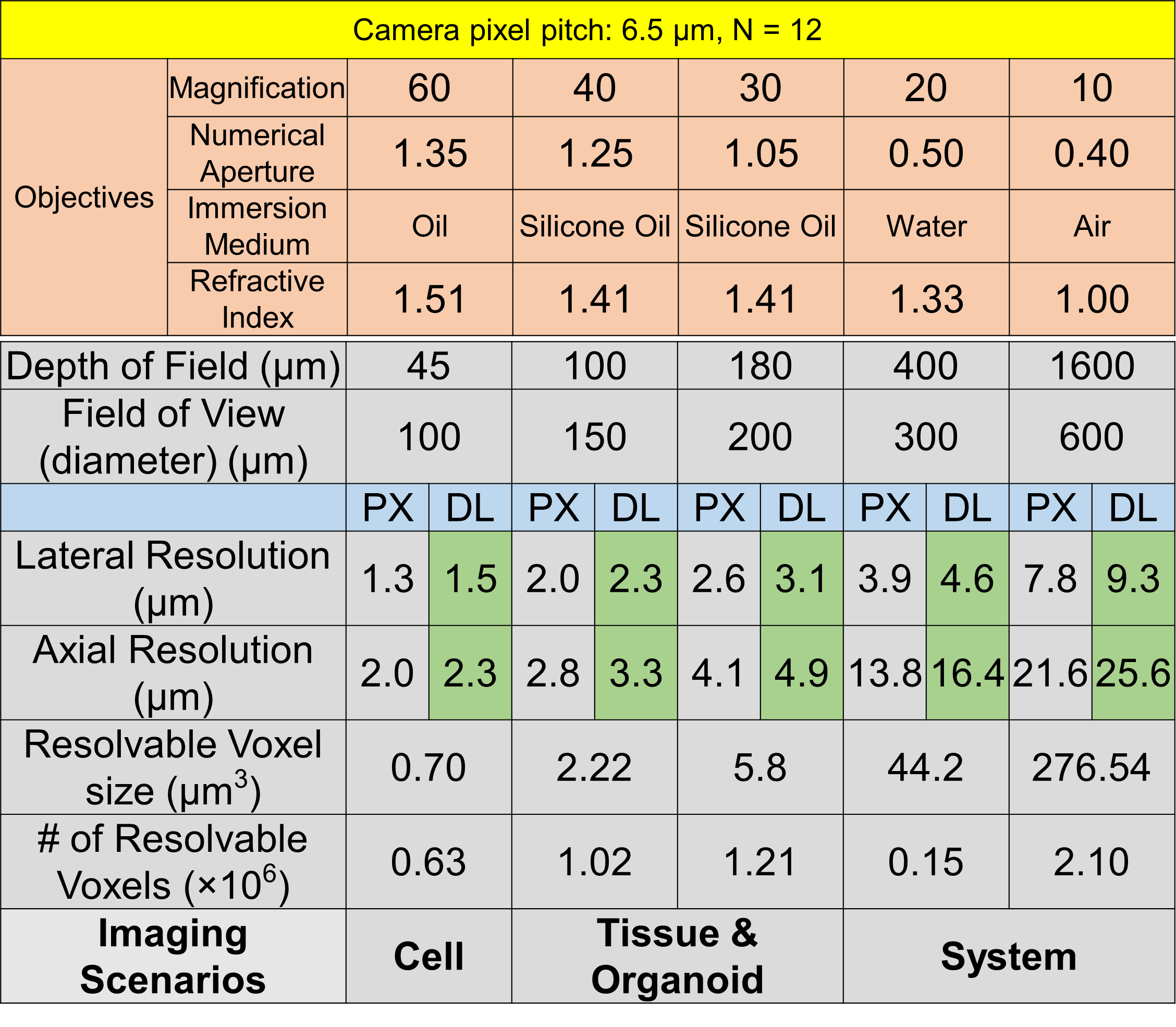

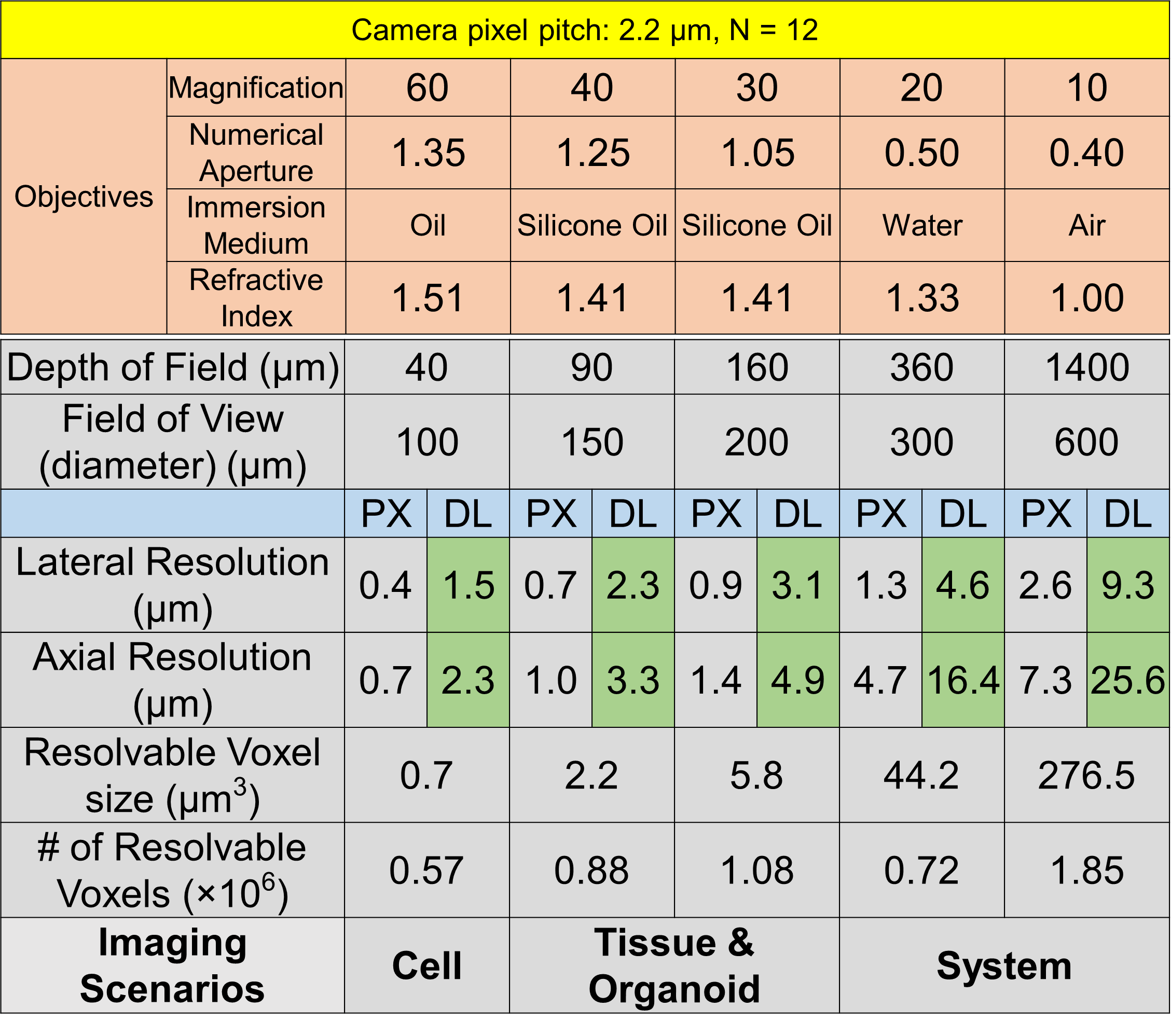

**Table 10**

**Table 9**

**Supplementary Table 11. The Number of Reconstructed Voxels vs the Number of Recorded Pixels.**

The Micro-lens pitch is 1.5 mm and the camera pixel pitch is 6.5 µm, which yields each circular micro-lens images with ~41500 pixels. The relay from the back aperture of the objective to the MLA is 1:1. Therefore, the size of illumination on to the MLA is the same as the size of the back aperture. The number (#) of microlens images is computed as: ${(\frac{Back Aperture Diameter}{1.5})}^{2}-4$, which is the number of micro-lenses illuminated. Here, “-4” represents the four corner micro-lenses that are not fully illuminated by the relayed aperture since the aperture has a circular shape. For all the objective of interest, there are more unknown (# of Voxels Reconstructed) than known variables (# of Pixels Recorded). Therefore, the 3D reconstruction is an ill-posed inverse problem for all the objectives considered. Comparing the ratio between the unknown vs the known, the 10X 0.4 NA objective poses a more well-determined system than the 60X 1.35 NA objective.

|  | 60X 1.35 NA  Oil | 40X 1.25 NA Silicone | 30X 1.05 NA Silicone | 20X 0.5 NA Water | 10X 0.4 NA  Air |
| --- | --- | --- | --- | --- | --- |
| Back Aperture Diameter (mm) | 5.3 | 7.9 | 8.9 | 6.7 | 14.4 |
| # of Microlens Images | 5  (3×3-4) | 12  (4×4-4) | 21  (5×5-4) | 12  (4×4-4) | 45  (7×7-4) |
| # of Pixels Recorded  (known) | 207500 | 498000 | 871500 | 498000 | 1867500 |
| # of Voxels Reconstructed  (unknown) | 846400 | 1745700 | 1957300 | 1375400 | 3332700 |
| $\frac{\mathbf{unknown}}{\mathbf{known}}$ | 4.1 | 3.5 | 2.1 | 2.8 | 1.8 |

**Supplementary Note 1. Calculation of performance parameters**

To calculate the imaging performance, we expressed the imaging performance parameters with respect to the optical parameters of the setup (Supplementary Table 2). The performance parameters are field of view (FOV), depth of view (DOV), lateral resolution ($R_{xy}$) and axial resolution ($R_{z}$). FOV, DOV and $R_{z}$ follow the same form derived in the Appendix of previous literature^1^. In the experiment, we translated laterally the MLA position for each objective such that the illuminated microlenses’ pattern is symmetrical. The theoretical performance estimate is calculated to align with this experimental procedure.

$R_{xy}$ is defined as the smallest resolvable distance between two point-sources. This distance is determined by two factors: the pixel pitch and diffraction limit of the LFM system. Based on Nyquist sampling theory, which states the sample rate must be greater than twice the highest frequency component of interest in the measured signal, the minimum distance that can be resolved by the camera pixel should be at least twice the distance of the pixel pitch. Mathematically, this is expressed as $R_{xy} \geq2p\cdot\frac{f_{2}}{{M\cdot f}_{\mathrm{ML}}}$ , ensuring that any signal variations over a distance greater than $R_{xy}$ can be sampled at least twice by the pixels. Additionally, the system's lateral resolution is constrained by the diffraction limit of light. Denote the ratio of relayed back aperture size at MLA ($ⅆ_{\mathrm{Rap}}$) over the pitch of each micro-lens $ⅆ_{\mathrm{ML}}$ as $N_{r}$ (Supplementary Fig. 4). Since each micro-lens is sub-sampling the main objective aperture, the image formed behind each micro-lens is a sub-aperture image, effectively reducing the numerical aperture (NA) by a factor of $N_{r}$. The diffraction limited resolution is expressed as $R_{xy} \geq\frac{\lambda N_{r}}{2NA}= \frac{\lambda}{2NA} \cdot\frac{2NA\cdot f_{2}}{M\cdotⅆ_{\mathrm{ML}}}=\frac{\lambda\cdot f_{2}}{M\cdotⅆ\_ML}$. The final lateral resolution is the larger one of these two distances.

The number of microlenses in the MLA is determined by leveraging the trade-off between the reconstructed single voxel size and total number of voxels, which were computed from equation (1) and (2) respectively for all the objectives of interest.

$voxel size= {(\frac{R_{xy}}{2})}^{2}\cdot\frac{R_{z}}{2}$	(1)

$\# of voxels= \frac{DOV\cdot\mathrm{FOV}^{2}}{voxel size}= \frac{8\cdot DOV\cdot\mathrm{FOV}^{2}}{{{R_{z\cdot}R}_{xy}}^{2}}$	(2)

To evaluate these two criteria with respect to N (the number of microlenses on one side of the square MLA), we can rewrite the equations in Supplementary Table 2 by substituting $ⅆ_{\mathrm{ML}}$ with N using equation (3), where $D_{MLA}$ is the diameter of the MLA.

$ⅆ_{\mathrm{ML}}= \frac{D_{MLA}}{N}$	(3)

The results were plotted in Figure 1b in the main text.

**Supplementary Note 2. Determination of number of microlenses in the MLA**

Under our system design with fixed f_2_ and f__ml_, the optimum number of microlenses can be achieved with an MLA with 9 x 9 microlenses. We chose an 8 x 8 MLA due to commercial availability. The final performance stayed close to optimum and satisfied a wide range of bio-imaging applications (Table 1). The main limiting factor on our LFM system is the pixel size. The camera we used was a standard scientific CMOS camera (Hamamatsu, ORCA-Flash4.0) with pixel size at 6.5 µm and pixel array dimension of 2048 × 2048. Reducing the camera pixel size will push the resolution to arrive at diffraction limit. Therefore, the optimum N will change accordingly.

As an example, with a camera of pixel size 2.2 µm, the performance improves with smaller N and the optimal choice for N would be at N = ~6 (Supplementary Fig 5). The imaging performance under this circumstance for three N’s are shown in Supplementary Table 6, 8, and 10. Compared to our camera with pixel size at 6.5 mm (Supplementary Table 5, 7, and 9), smaller pixel size allows for higher pixel sampling rate and therefore, the imaging performance prefers applications with higher imaging resolution and smaller DOV. However, smaller pixel size sometimes comes with lower quantum efficiency and lower frame rate. The final choice could be determined with a combination of imaging requirements, cost of manufacturing and commercial availability. Our contribution is focused on providing a metric and methodology with open-source design software to help people to calculate and decide on different hardware parameters for their own experiments.

**Supplementary Note 3. Experimental PSF processing and 3D reconstruction of LFM data.**

The complete processing pipeline, from experimental PSF extraction to 3D reconstruction, was implemented in MATLAB and FIJI. It is open-source and publicly available on GitHub with a detailed step-by-step manual that does not require expert knowledge of LFM [*https://github.com/yangyanb/FLFM-Processing-Pipeline.git*]. FIJI was used for basic image pre-processing including background subtraction and generating binary masks for regions of interest (ROI). MATLAB scripts were written for extracting the experimental PSF from the beads’ image stack, cropping out the ROI in the raw LFM image then performing 3D reconstruction. The 3D reconstruction section was adapted from a previous package named *oLaF*^2–4^. The reconstructed 3D data were generated in both TIF and MAT format for immediate visualization and downstream processing.

The pipeline consists of two processing stages: experimental PSF extraction and 3D reconstruction. The experimental PSF was collected from scanning a single bead axially over the DOV placed in the center of the FOV. The bead image at different axial planes shifts its center positions laterally in each angular projection view (Fig. 2a, b, Supplementary Fig. 2a, b, d, e). For this reason, a direct cylindrical cut-out across the entire bead image stack is not ideal since it includes large unrelated background volume that contains pure background shot noise and could contain contamination caused by other beads or fluorescent objects, such as dust and bubbles. A proper PSF extraction process ensures a clean and accurate PSF characterization of the system, reducing computational artifacts in downstream 3D reconstruction. Here, we develop a PSF extraction pipeline that is specifically tailored to the LFM system. Given that the shift between adjacent axial slice does not change dramatically (around 1-2 pixels), the bead region in the current slice would still be contained in the masked region of the last slice. This property was exploited to iteratively update the PSF mask at every plane so that the mask is always positioned near the center of the bead image in each angular projection view. To initiate the iteration, the bead image stack was circled manually in the first axial plane. The iteration is written in MATLAB scripts and described in the following steps:

***Step 1:*** Subtract background from the bead stack.

***Step 2:*** Circle out Regions of Interest (ROIs) for a single bead in all angular projection views manually at the 1st slice of the bead stack.

***Step 3:***

- Create a bead mask from the ROIs.
- Mask out the 1st slice of the bead stack.

***Use i to denote the slice number, then from i = 2 to the last slice:***

***Step 4:***

- Read in the i^th^ slice.
- Find the maximum intensity position in each masked-out region.

***Step 5:***

- Shift the bead mask at i-1^th^ slice in each angular projection view such that the center is the maximum intensity position in Step 3.
- Use the shifted mask as the bead image mask for i^th^ slice.

***Step 6:***

- Mask out the single bead light field image at i^th^ slice.
- i = i + 1.

The 3D reconstruction algorithm takes the extracted PSF and the raw light field image to iteratively reconstruct the 3D volume. As shown in the method section, the main operation in the iterations is multiple 2D convolutions for forward and backward projection between the 2D light field image space and 3D sample space. Graphics Processing Unit (GPU) computing was used to accelerate the iterations. We used one Nvidia Quadro p5000 GPU with 16 GB of RAM on a workstation for all the reconstructions. With a fix number of iteration of 10, the reconstruction time varied depending on the size of the light field images. The reconstruction time for each experiment is listed in Supplementary Table 3. Roughly speaking, the reconstruction time is at ~ 30 to 600 folds of the acquisition time, varying on the size of the light field image data as well as the reconstructed number of z slices. To speed up the reconstruction, we could improve the hardware computing power as well as the software implementation. On the hardware side, apart from increasing the GPU processing speed and RAM volume, Cloud Computing Service offered by companies such as Amazon offers a cost-effective way to boost the data processing power for reconstructing large datasets. On the software side, since the reconstruction time goes down with decreasing the size of the raw 2D LFM image, down sampling the light field image before reconstruction can shorten the reconstruction time significantly (Supplementary Fig. 3). While this approach inevitably compromises resolution, it is often useful when the resolution exceeds the necessary requirements for imaging dynamics, producing redundant data that offers little additional insight.

The reconstruction problem is formulated as an inverse problem. The robustness of the reconstruction is determined by how well-pose the inverse problem is under different scenarios. For our setup, the 10X 0.4 NA objective yields a greater number of angular views, i.e. microlens images (7×7 array, 45 in total) compared to other objectives (Supplementary Table 11). However, as illustrated in Figure 1c of the Main Text, the total number of resolvable voxels—the unknowns to be estimated in the inverse problem—is also higher for the 10X 0.4 NA objective. Therefore, a higher number of views does not necessarily lead to a more robust or over-determined inverse problem. The robustness of the inverse problem depends not only on the number of angular views, but also the spatial-angular information captured by the system. This is influenced by several key factors:

1. The native numerical aperture (NA) of the objective lens.
2. The number of microlenses used for reconstruction. which controls angular sampling density.
3. The number of camera pixels behind each microlens, which governs the spatial sampling within each view.

It is the balance and interaction among these factors—not the number of views alone—that determines the conditioning and stability of the inverse problem. A detailed comparison of the number of unknown voxels reconstructed versus recorded pixels is listed in **Supplementary Table 11**. As seen from the Table, under our system design, the 10X 0.4 NA objective does pose a more well-determined system than rest of the objectives.
